# Balancing the length of distal tip is key for stability and signalling function of primary cilia

**DOI:** 10.1101/2021.06.04.447095

**Authors:** Taishi Kanamaru, Annett Neuner, Bahtiyar Kurtulmus, Gislene Pereira

## Abstract

Primary cilia are antenna-like organelles required for signalling transduction. How cilia structure is mechanistically maintained at steady-state to promote signalling is largely unknown. Here, we define that mammalian primary cilia are formed by middle and distal segments, in analogy to sensory cilia of lower eukaryotes. The analysis of middle/distal segmentation indicated that perturbations leading to cilia over-elongation influenced middle or distal segment length with a different impact on cilia behaviour. We identified Septins as novel repressors of distal segment growth. We show that Septins control the localisation of MKS3 and CEP290 required for a functional transition zone, and through this the entrance of the microtubule-capping kinesin KIF7, a cilia-growth inhibitor, into the cilium. Live-cell imaging and analysis of sonic-hedgehog (SHH) signalling activation established that distal segment over-extension increased cilia excision events and decreased SHH activation. Our data underlies the importance of understanding cilia segmentation for length control and cilia-dependent signalling.

## Introduction

Primary cilia are conserved, specialized antenna-like organelles that protrude from the cell surface where they perceive signals from the extracellular milieu and, as a consequence, activate downstream responses to the cell interior (Ishikawa & Marshall, 2011; Wheway *et al*, 2018; Lee *et al*, 2015). As such, the primary cilium plays essential roles during animal development and tissue homeostasis through regulation of important signalling pathways such as Wnt, Hedgehog (HH) and TGF1-β (Echelard *et al*, 1993; Wheway *et al*, 2018; Mönnich *et al*, 2018; May-Simera *et al*, 2018). Therefore, it is not surprising that defective cilia can lead to multi-organ dysfunction due to signalling failure, which is the aetiology of genetic disorders named ciliopathies (Leightner *et al*, 2013; Slaats *et al*, 2016; Waters & Beales, 2011; Guen *et al*, 2016; Reiter & Leroux, 2017).

The cilium is built from the basal body, through a cascade of events that lead to transition zone (TZ) establishment, cilia membrane formation and axoneme microtubule (axoMTs) extension (Ishikawa & Marshall, 2011; Sánchez & Dynlacht, 2016; Kim & Dynlacht, 2013). Once formed, cilia are maintained at a specific length that varies between cell types (Besschetnova *et al*, 2010; Wann & Knight, 2012; Qiu *et al*, 2012; Miyoshi *et al*, 2014; He *et al*, 2014; Ghossoub *et al*, 2013). For instance, relatively long cilia are reported in neurons (8-10µm), whilst much shorter cilia are present in chondrocytes (2µm) (Wann & Knight, 2012; Miyoshi *et al*, 2014). Variations in the physiological length of cilia causing shortening or over-elongation are reported in ciliopathies, indicated that keeping proper cilia length in each tissue is vital for their signalling function (Guen *et al*, 2016; Ramsbottom *et al*, 2018; Leightner *et al*, 2013; May-Simera *et al*, 2018; Reiter & Leroux, 2017; Kim *et al*, 2018).

Evidence indicates that ciliary organization as well as identity is not uniform. The structure of the axoneme has been studied in *C. elegans* sensory cilia where it has been sub-divided into two domains, named the middle (MS) and distal segments (DS) (Hao *et al*, 2011; Kramer *et al*, 2007). The MS is defined by the doublet MTs, which extend from the basal body and are enriched in several post-translational modifications that stabilise axoMTs, such as polyglutamylation and polyglycylation (Snow *et al*, 2004b; O’Hagan *et al*, 2011; Kramer *et al*, 2007; Alford *et al*, 2017). The DS defines the domain in which doublet MTs converge into singlets, which extend from the A-tubule of axoMTs towards the cilia tip (Hao *et al*, 2011; Kramer *et al*, 2007; van der Burght *et al*, 2020; Mukhopadhyay *et al*, 2008). The DS is proposed to act as a chemical and osmotic sensor (Kramer *et al*, 2007; Alford *et al*, 2017; van der Burght *et al*, 2020; Cornelia I. Bargmann, 2006). Recent electron tomography data shows that the doublet-singlet MT transition also occurs in mouse epithelial primary cilia (Sun *et al*, 2019; Flood & Totland, 1977) and signalling pathway-related proteins are enriched at the ciliary tip, reinforcing the importance of the DS in signalling (Pedersen & Akhmanova, 2014; Haycraft *et al*, 2005; Wheway *et al*, 2018; Cherry *et al*, 2013). Therefore, it is strongly indicated that this middle-distal segmentation is structurally conserved in mammalian cilia. However, it is still unclear how these segments influence primary cilia function in mammalian cells.

Here, our data show that human and mouse primary cilia contain middle-distal segments, separated by enrichment of polyglutamylated tubulin. Interestingly, factors that led to cilia over-extension impact the size of the middle-distal segments differently. We identify Septin as a novel factor that specifically represses DS extension, without changing the MS. Septin controls localisation of TZ components MKS3 and CEP290 to transfer the axonemal-capping kinesin KIF7 to the ciliary distal tip for DS growth. Moreover, DS hyper-elongation induces cilia excision due to structural instability and causes defective SHH signalling activation. We propose that the control of the DS by Septin is key for the maintenance of primary cilia signalling function. Our data underlies the importance of understanding how cilia sub-domains are maintained and regulated.

## Results

### Observation of cilia segmentation in human and mouse cells

To understand the mechanisms controlling cilia length, we characterized the dynamics of cilia elongation in human retinal pigment epithelial (RPE1) and mouse fibroblast (NIH3T3) cells after serum-starvation, a condition that induces cilia biogenesis in a timely manner (Fig. 1a-c, S1a-c). In both cell types, cilia reached maximum length after 24h serum-starvation which was maintained for 48h (Fig. 1b, S1b). To better characterize which part of the axoneme contributed to cilia growth, we determined the middle-distal segments in mammalian cells, as reported for *C. elegans* (Lechtreck & Geimer, 2000; Snow *et al*, 2004b; Cornelia I. Bargmann, 2006; Hao *et al*, 2011). For this, we stained for axoMT markers, acetylated and polyglutamylated-tubulin, which stain singlet/doublet and doublet MTs respectively (He *et al*, 2020), and the membrane marker ARL13B (Fig. 1d). The stained regions of cilia were similar when comparing ARL13B and acetylated tubulin staining using conventional or stimulated emission depleted microscopy (STED) in RPE1 (Fig. 1e, S1d-e). This pattern was, however, different for glutamylated tubulin, which strongly decorated the middle part of the cilium (Fig. 1d, S1d). Based on these data, we defined the MS as the polyglutamylated-tubulin rich region, whilst the length of the DS was calculated as the difference between the full cilium (ARL13B-stained region) and the MS (Fig. 1d). Interestingly, shorter cilia (< 2 µm) had a very short DS, but in longer cilia (> 4 µm) the DS represented more than 30% of the cilium in human and mouse cells (Fig. 1f, S1f). In both cell types, longer DS were more predominant after 48h of serum-starvation (Fig. 1g, S1g). Together, these data imply that cilia growth involves timely elongation of the MS followed by the DS.

**Figure 1.**
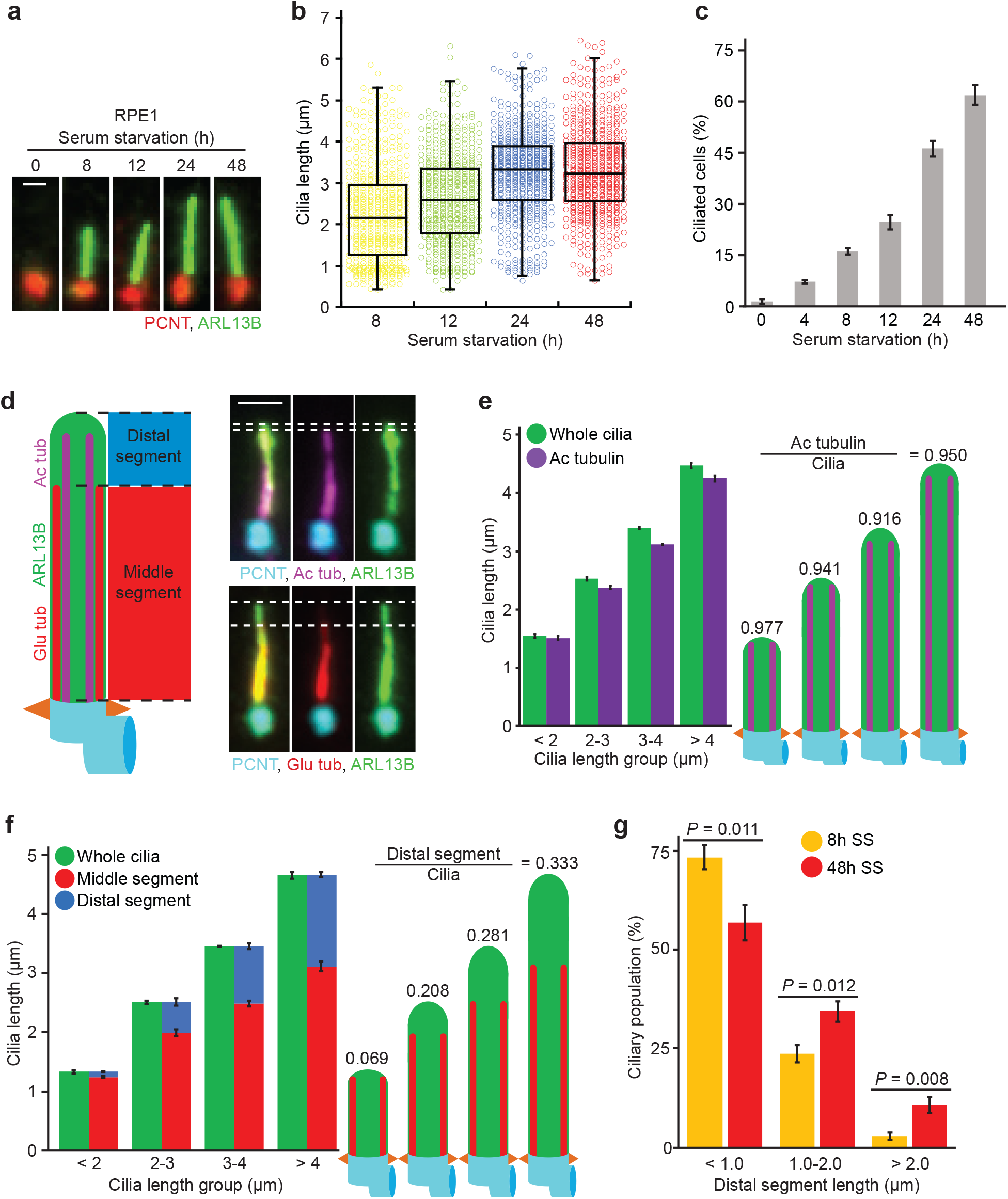
Characterisation of middle/distal cilia segmentation in RPE1 cells. **a-b**, Representative images of cilia (**a**) and dot plots (**b**) showing primary cilia length variation at different time points after serum starvation (SS). Markers in (**a**) show ARL13B (cilia) and PCNT (Basal body). The graph shows individual length of cilia from 3 biological replicates (n>150 cilia per sample and experiment). Scale bar: 1µm. **c**, Percentage of ciliated RPE1 cells after SS from **a**. 3 biological replicates, n>100 cells per sample and experiment repetition. **d**, Representative images show cilia segmentation based on acetylated tubulin (Ac tub), polyglutamylated tubulin (Glu tub), ARL13B and PCNT. The cartoon on the left indicates middle segments (MS, Glu tub rich-region) and distal segments (DS). DS were determined by calculating the difference between MS and total cilia length based on ARL13B staining as depicted by the dashed lines on the images. Scale bar: 1.5µm. **e**, Quantification of **d**. Comparative analysis of average cilia length based on Ac tubulin and ARL13B (cilia) staining shown for specified cilia length groups (2<, 2-3, 3-4 and >4µm). Three biological replicates, n=400 cilia per sample and repetition. The numbers above the cartoons show the ratio of the calculated Ac tub/whole cilia length. **f**, Average length of MS and DS for the cilia length groups depicted. Representative images (**d**) and quantifications of MS (Glu tub) and whole cilia (ARL13B) length are shown. The numbers above the cartoons show the ratio of the calculated DS/whole cilia length. Three biological replicates, n>650 cilia per sample and repetition. **g**, Percentage of cilia with depicted DS length after 8h and 48h SS. Three biological replicates, n>150 cilia per sample and repetition. Data shown in (**b, c, e, f** and **g**) include mean ±s.d. and *P* values of (**g**) are calculated by two-tailed unpaired student t-test. Source data: Supplementary Table 1.

### Microtubules and Septins affect DS growth

We next evaluated how the MS and DS change their length when using low-doses of nocodazole or cytochalasin D (CytoD), which were reported to elongate cilia without collapsing the MT and actin cytoskeletons, respectively(Sharma *et al*, 2011; Kim *et al*, 2010). Treatment of RPE1 and NIH3T3 cells with nocodazole or CytoD led to a significant increase in cilia extension analysed by acetylated tubulin and/or ARL13B (Fig. 2a-b, Fig. S2a-c). CytoD elongated both MS and DS length without changing the segment ratio, whereas nocodazole increased only the DS (Fig. 2a-b, S2a-c). Dual nocodazole and CytoD treatment additionally extended DS length in comparison to CytoD alone in RPE1 cells (Fig. 2a-b). Together, MT and actin perturbations increase cilia length by affecting different segments.

**Figure 2.**
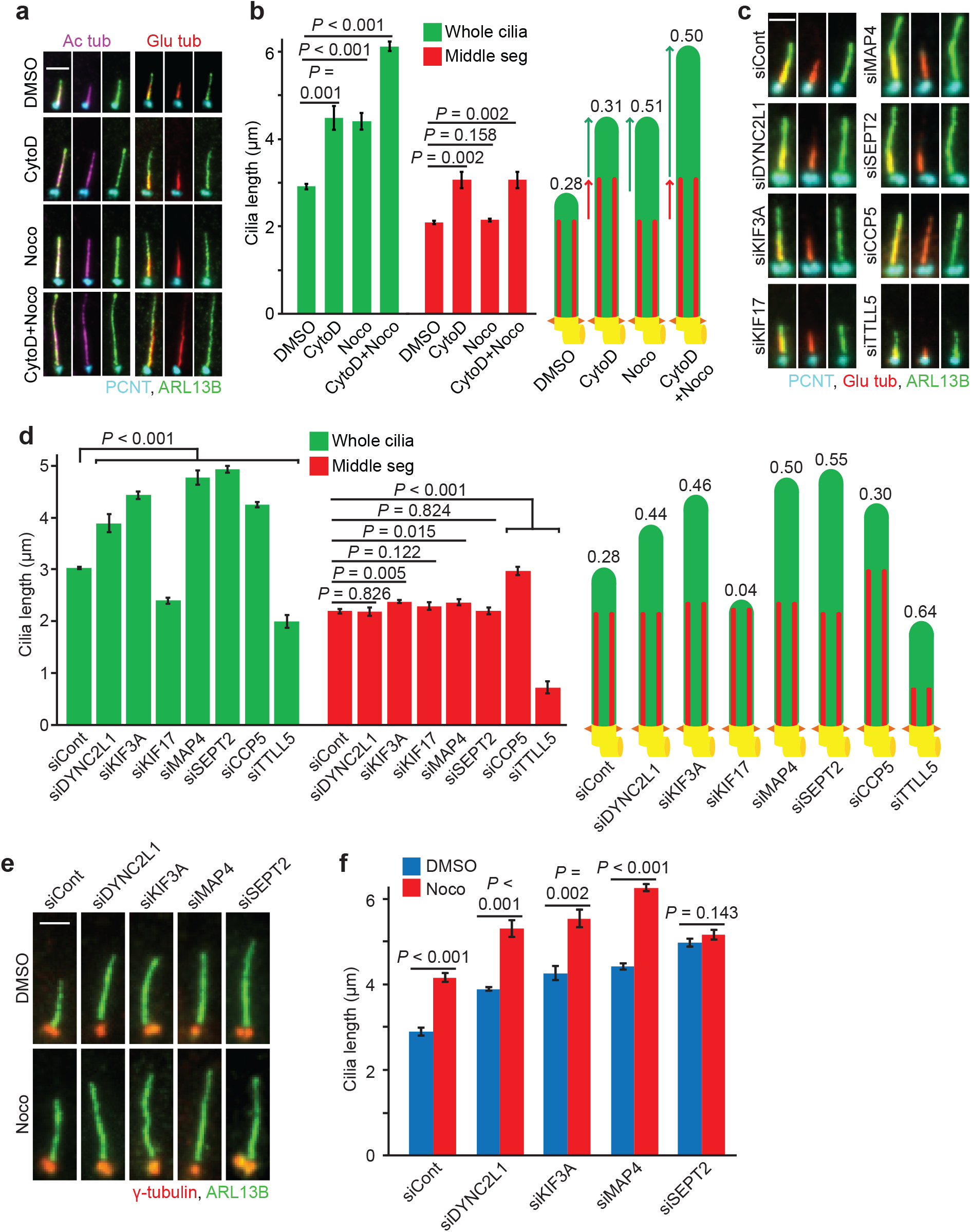
Microtubule perturbators and SEPT2 depletion change only the distal segment length in RPE1 cells. **a,** Representative images of 48h serum starved RPE1 cells treated with solvent control (DMSO), cytochalasin D (CytoD) and/or nocodazole (Noco) for 3h before immunostaining with the indicated antibodies. Scale bar: 3µm. **b,** Quantification of (**a**) showing average length of middle segment (MS, Glu tub) and whole cilia (ARL13B). The numbers above the cartoons show the ratio of the calculated distal segment (DS)/whole cilia length. Three biological replicates, n=100 cilia per sample and repetition. **c-d**, RPE1 cells were treated with the indicated siRNAs before serum starvation for 48h. Representative immunostained images with indicated antibodies (**c**) and quantifications of MS (Glu tub) and whole cilia (ARL13B) length (**d**) are shown. The numbers above the cartoons in (**d**) show the ratio of the calculated DS/whole cilia length. Scale bar: 2µm. Three biological replicates, n=100 cilia per sample and repetition. **e-f**, Cells were treated with the indicated siRNAs before serum starvation for 48h and treatment with DMSO or Noco for 3h before immunostaining with the indicated antibodies. Representative images (**e**) and quantification of cilia length based on ARL13B (**f**) are shown. Scale bar: 2µm. Three biological replicates, n=100 cilia per sample and repetition. Data shown in (**b**, **d** and **f**) include mean ±s.d.; *P* values are calculated by two-tailed unpaired student t-test. Source data: Supplementary Table 1.

We next became interested in identifying proteins that control DS elongation upon nocodazole treatment. We reasoned that depletion of these proteins should lead to elongated cilia by causing only DS and not MS extension. Additionally, depletion of these candidates should not further increase cilia length upon nocodazole treatment. Using these criteria, we tested several candidate genes in an siRNA-based screen. The encoded proteins were reported to either interact with MTs or affect cilia length and included the IFT dynein-2 light-chain DYNC2L1 (Taylor *et al*, 2015), IFT kinesin-subunits KIF3A (Qiu *et al*, 2012) and KIF17 (Insinna *et al*, 2008), MT-interacting proteins MAP4 and SEPT2 (Hu *et al*, 2010; Ghossoub *et al*, 2013), tubulin de-glutaminase CCP5 (He *et al*, 2018) and glutaminase TTLL5 (Wloga *et al*, 2017; Sun *et al*, 2016). The knockdown (KD) of these genes, excluding KIF17 and TTLL5, increased ciliary length in comparison to the control (Fig. 2c-d). CCP5-KD extended both segments, keeping segment proportion as the control (Fig. 2d). Similar to nocodazole treatment, DS length was raised upon DYNC2L1, KIF3A, MAP4 and SEPT2-KD (Fig. 2d). Of those, nocodazole treatment further extended cilia length in DYNC2L1, KIF3A and MAP4 but not SEPT2-KD cells (Fig. 2e-f). We thus hypothesized that SEPT2 and MTs might be involved in the same pathway regulating DS elongation and decided to investigate this function of Septins in more detail.

### Septins are enriched along the cilia but not at the TZ

SEPT2 is one of the building blocks of the Septin family, which is sub-divided into four sub-groups represented by SEPT2, SEPT6, SEPT7 and SEPT9(Neubauer & Zieger, 2017). Septins support cell division and act as membrane diffusion barriers in different contexts and organisms (Sandrock *et al*, 2011; Valadares *et al*, 2017; Palander *et al*, 2017; Ghossoub *et al*, 2013). In RPE1 cells, Septins were reported to form filamentous structures in the cytoplasm and cilium (Ghossoub *et al*, 2013). Indeed, specific antibodies against SEPT2 and SEPT7 confirmed this localisation (Fig. 3a). We then asked whether Septins decorate the entire cilium including the TZ. For this, we used a functional EGFP-SEPT2 fusion protein (Fig. S3) for co-localisation with MKS3 or CEP290 as TZ markers (Garcia-Gonzalo & Reiter, 2017; Shi *et al*, 2017) alongside ARL13B and γ-tubulin (Fig. 3b). The lengthwise comparison of the signal intensities showed that EGFP-SEPT2 signal was very low at MKS3 or CEP290 regions but increased along the cilia and then slightly diminished towards the tip (Fig. 3c). This indicates that Septins accumulate along the cilium but not at the TZ.

**Figure 3.**
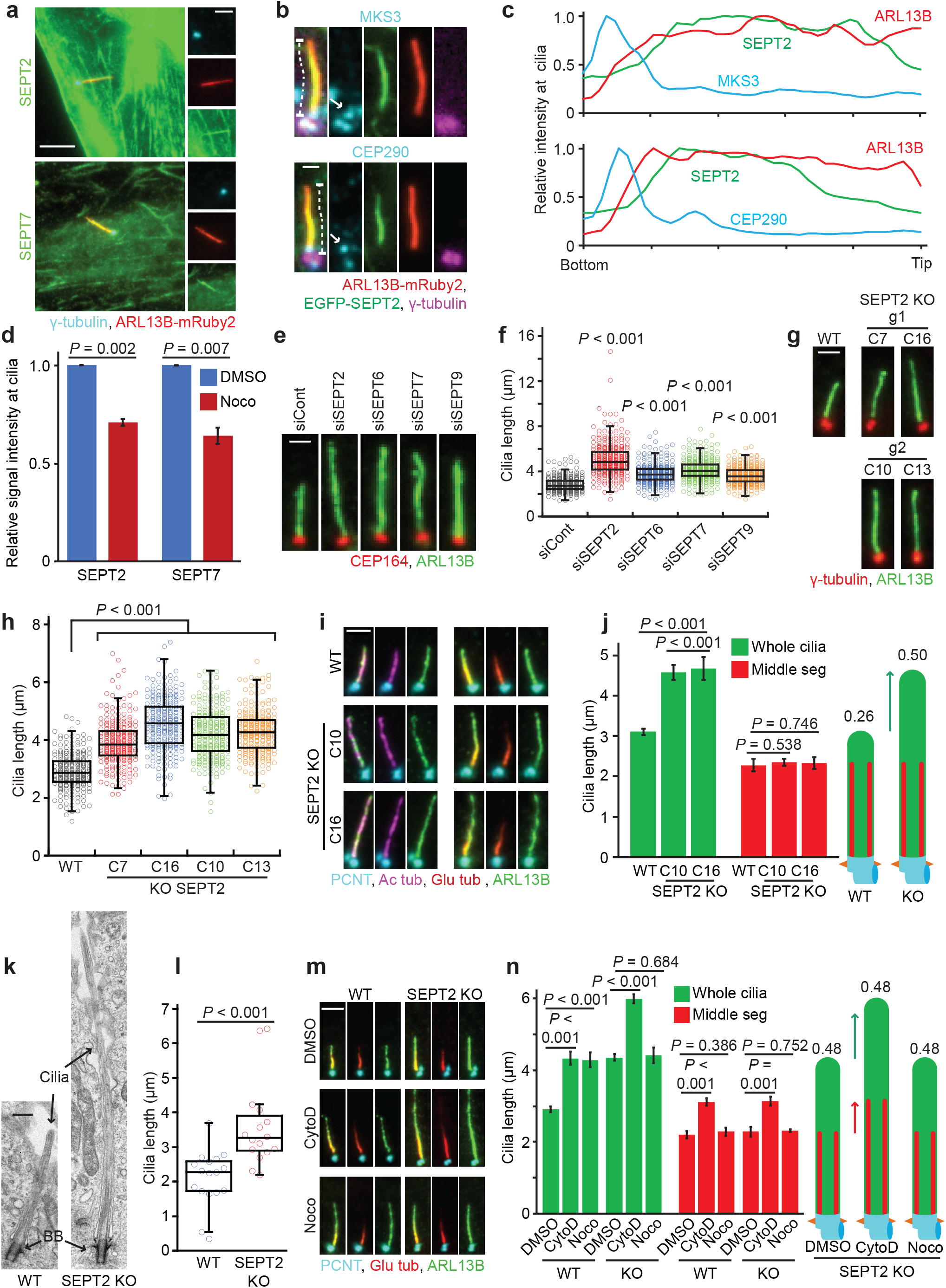
Septin negatively controls distal segment elongation in RPE1 cells. **a,** Micrographs show the localisation of SEPT2 and SEPT7 in RPE1 ARLB13B-mRuby2 cells with basal body staining. The insets to the right show enlargements of the cilium. Scale bar: 4µm (large) and 2µm (small). **b**, Representative images showing co-localisation of GFP-SEPT2 and MKS3 or CEP290 in ciliated RPE1 ARLB13B-mRuby2 cells. The white arrows point to the specific MKS3 or CEP290 signals. Scale bar: 1µm. **c,** Quantification of **(b).** Relative signal intensities were determined along the cilium for the indicated proteins as specified in (**b**) (dashed lines). **d,** Relative signal intensity of endogenous SEPT2 or SEPT7 along the cilium in cells treated with solvent control (DMSO) or nocodazole (Noco) for 3h after 48h serum starvation (SS). SEPT2 or SEPT7 signals at cilia were normalized to the total cilia length. Three biological replicates, n>90 cilia per sample and repetition. See Fig. S3a for representative images. **e-f,** Cilia length changes in RPE1 cells treated with the indicated siRNAs and 48h SS stained with indicated antibodies. Representative images of cilia (**e**) and quantifications (**f**) are shown. Dot plots in (**f**) represent individual cilia. Scale bar: 1.5µm. Three biological replicates, n=100 cilia per sample and repetition. **g-h,** Cilia length changes in RPE1 SEPT2-KO single clones after 48h SS stained with indicated antibodies. Representative images of cilia (**g**) and quantifications (**h**) are shown. Dot plots in (**h**) represent individual cilia. The parental cells (WT) and four KO clones (C7, C16, C10, C13) derived from two independent gRNAs (g1 and g2, as depicted in **g**) were analysed. Three biological replicates, n>80 cilia per sample and repetition. Scale bar: 2µm. **i-j,** Cilia segmentation in SEPT2-KO ciliated cells serum starved for 48h. Cells were immunostained with the indicated antibodies. Representative images (**i**) and quantifications of middle segment (MS, Glu tub) and whole cilia (ARL13B) length (**j**) are shown. The numbers above the cartoons in (**j**) show the ratio of the calculated distal segment (DS)/whole cilia length. Scale bar: 2µm. Three biological replicates, n=100 cilia per sample and repetition. **k,** Electron microscopy of RPE1 WT and SEPT2-KO cilia. Black arrows point to the axoneme and basal body (BB). Scale bar: 400nm. **l,** Quantification of **k.** Only cilia detected from bottom (BB) to tip in serial sections were quantified. N>15 cilia from two independent experiments. **m-n**, RPE1 WT and SEPT2-KO cells after 48h SS were treated with solvent control (DMSO), cytochalasin D (CytoD) and/or nocodazole (Noco) for 3h before immunostaining with the indicated antibodies. Representative images (**m**) and quantifications of MS (Glu tub) and whole cilia (ARL13B) length (**n**) are shown. The numbers above the cartoons in (**n**) show the ratio of the calculated DS/whole cilia length. Scale bar: 3µm. Three biological replicates, n=100 cilia per sample and repetition. Data show in (**d**, **f**, **h**, **j**, **l**) and **n** include mean ±s.d.; *P* values are calculated by two-tailed unpaired student t-test (in **d, j, n**) or by unpaired Wilcoxon-Mann-Whitney Rank Sum Test (in **f, h, l**). Source data: Supplementary Table 1.

As nocodazole treatment increased DS extension similar to SEPT2-KD, we asked whether Septin cilia localisation could be influenced by nocodazole. Interestingly, nocodazole treatment significantly reduced endogenous SEPT2 and 7 levels at the cilium in RPE1-WT cells (Fig. 3d, S3a), indicating that Septin cilia localisation is partly influenced by the MT cytoskeleton.

### Septins suppress DS extension

To test whether the effect on cilia elongation was limited to SEPT2, we compared cilia length upon depletion of SEPT2, SEPT6, SEPT7 and SEPT9 in RPE1 cells and in all cases cilia were longer (Fig. 3e-f, S3b). Contra to a previous study(Ghossoub *et al*, 2013), Septin-depleted cells were able to ciliate more efficiently than control-depleted cells (Fig. S3c). To exclude off-target effects, we generated doxycycline (DOX) inducible constructs carrying siRNA-resistant EGFP-SEPT2, EGFP-SEPT6 and SEPT7-EGFP. EGFP-SEPT6 localised along cilia similarly to SEPT2 and SEPT7 (Fig. S3d). The rescue constructs reverted cilia length in SEPT2, 6 and 7 depleted cells (Fig. S3e), excluding off-target effects.

Next, we constructed RPE1 SEPT2 knockout (KO) cells using a CRISPR/Cas9 approach. Using two different guide-RNAs (gRNA), we obtained four distinct SEPT2-KO clones (clones 7, 10, 13 and 16) that completely lost endogenous SEPT2 (Fig. S3f-g). All SEPT2-KO cells produced longer cilia and showed higher ciliation rates compared to WT cells (Fig. 3g-h, S3h). Cilia over-elongation was rescued by EGFP-SEPT2 (Fig. S3i). Similar to siRNA depletion, only the DS was elongated in SEPT2-KO cells (Fig. 3i-j, S3j). Longer cilia in SEPT2-KO cells were also observed by electron microscopy (Fig. 3k-l). Additionally, the treatment of SEPT2-KO cells with nocodazole had no further effect on DS extension, whereas CytoD treatment elongated the DS and MS (Fig. 3m-n). Similar to RPE1 cells, SEPT2 and SEPT7-KD in NIH3T3 cells formed longer DS without any effect on ciliogenesis (Fig. S3k-n). Nocodazole treatment in NIH3T3 SEPT2-KD cells did not promote further DS elongation in comparison to the control (Fig. S3o-p). Together, these data strongly indicate that Septins are involved in length control of the ciliary DS in human and mouse cells.

### Localisation of the ciliary-capping protein KIF7 at cilia tip requires Septin

We reasoned that the elongated DS phenotype in Septin loss could be a consequence of imbalanced localisation of promoters or inhibitors of cilia growth. One key factor is the IFT complex machinery, which works to bi-directional transport components in and out of cilia (Snow *et al*, 2004a; Prevo *et al*, 2017). We thus compared the amount of two IFT components, IFT88 and KIF17, at the cilium. The signal intensities along the axoneme showed no significant difference between RPE1-WT and SEPT2-KO cells (Fig. 4a-b), suggesting that Septins do not affect the localisation of IFT complexes along cilia.

**Figure 4.**
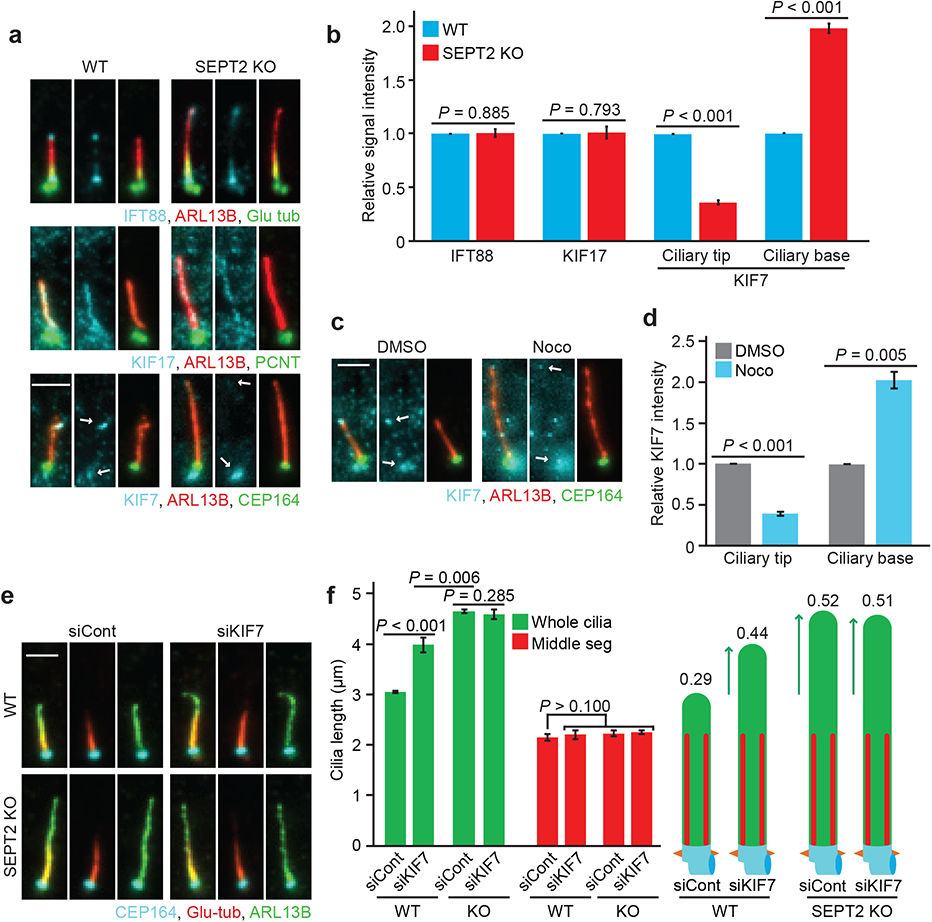
Septin is required for the KIF7 accumulation at the cilia tip to suppress distal segment growth. **a-b**, Representative images show the localisation of IFT88, KIF17 and KIF7 in RPE1 WT and SEPT2-KO cells after 48h serum starvation (SS) stained with indicated antibodies (**a**) and quantification of the relative signal intensities for the indicated proteins (**b**). White arrows point to the base and tip localisation of KIF7. Scale bar: 3µm. IFT88 and KIF17 signal intensities were normalized to the whole cilia length. KIF7 signal intensity was quantified at ciliary tip or base as indicated in (**a**) (white arrows). Three biological replicates, n>80 cilia per sample and repetition. **c-d**, KIF7 localisation at the tip and base of cilia (indicated by arrows) in RPE1 cells after 48h SS treated for 3h with solvent control (DMSO) or nocodazole (Noco) and quantification of the relative signal intensities for the ciliary base and tip of KIF7 (**d**). Scale bar: 2µm. KIF7 signal intensity was quantified at the ciliary tip or base as indicated in (**c**) (white arrows). Three biological replicates, at least 80 cilia per sample and repetition. **e-f**, RPE1 WT and SEPT2-KO cells were treated with control or KIF7 siRNA after 48h SS and immunostained with the indicated antibodies. Representative images (**e**) and quantifications of middle segment (MS, Glu tub) and whole cilia (ARL13B) length (**f**) are shown. The numbers above the cartoons in (**f**) show the ratio of the calculated DS/whole cilia length. Three biological replicates, n=100 cilia per sample and repetition. Scale bar: 2µm. Data shown in (**b**, **d**, **f**) include mean ±s.d. and *P* values are calculated by two-tailed unpaired student t-test. Source data: Supplementary Table 1.

Next, we investigated the axoMT-capping kinesin KIF7. KIF7 is a repressor of cilia elongation that binds to the distal end of axoMTs to stop their growth (He *et al*, 2014; Pedersen & Akhmanova, 2014; Lewis *et al*, 2017). KIF7 localised at the cilia base and tip in RPE1 cells (Fig. 4a). In SEPT2-KO cells, KIF7 levels significantly decreased at the ciliary tip, whilst it accumulated at the ciliary base (Fig. 4a-b). EGFP-SEPT2 expression in SEPT2-KO cells recovered KIF7 localisation at the ciliary tip (Fig. S4a-b). The specificity of KIF7 localisation at cilia in RPE1 cells was proven using KIF7-KD (Fig. S4c-d). KIF7 at the ciliary tip was also reduced in NIH3T3 cells upon KIF7, SEPT2 and SEPT7-depletion (Fig. S4e-f). Furthermore, KIF7 mis-localisation was phenocopied in nocodazole-treated RPE1 and NIH3T3 cells (Fig. 4c-d, S4g-h), suggesting that KIF7 might be the underlying mechanism for DS over-elongation. To test this, we analysed KIF7-KD in RPE1 and NIH3T3 cells. KIF7-depletion did not alter ciliogenesis but extended the DS (Fig. 4e-f, S4i-l). Remarkably, KIF7 depletion in RPE1 SEPT2-KO cells or SEPT2/KIF7 or SEPT7/KIF7 co-depletion in NIH3T3 cells did not cause further DS elongation (Fig. 4e-f, S4k-l). Together, these data strongly indicate that Septins are required for KIF7 entry into the cilium, where it controls DS length. It also implies that Septins, MTs and KIF7 most likely are part of the same pathway controlling DS elongation, with KIF7 functioning downstream of Septins.

### MKS3 and CEP290 require SEPT2 for their proper localisation at the TZ in RPE1 cells

Considering that the TZ acts as a gate-keeper to selectively control entry of ciliary components into the cilium (Garcia-Gonzalo & Reiter, 2017; Lee *et al*, 2015), we asked whether it could be impaired in cells lacking Septins. We investigated the localisation of TZ components MKS3, CEP290 and NPHP1 (Garcia-Gonzalo & Reiter, 2017). In RPE1 cells, the specific MKS3 signal co-localised with CEP164 at the ciliary base (Fig. 5a, arrow), whilst a non-specific signal was detected close to the basal body (Fig. 5a, asterisk), as this signal did not disappear upon MKS3 depletion (Fig. S5a). In SEPT2-KO cells, the specific MKS3 signal dramatically extended from the base into cilia (Fig. 5a-b). STED microscopy confirmed MKS3 mis-localisation along the cilium in SEPT2-KO cells (Fig. S5c). CEP290 or NPHP1 did not extend along cilia (Fig. 5a), however, the levels of CEP290 at the TZ significantly increased in SEPT2-KO cells (Fig. 5a, c). MKS3 and CEP290 mis-localisation were recovered by EGFP-SEPT2 expression in SEPT2-KO cells (Fig. S5d-f). CEP164 and ODF2 levels remained unaltered in SEPT2-KO cells, suggesting that SEPT2-KO does not affect basal body formation (Fig. 5c). Furthermore, nocodazole treatment phenocopied SEPT2-KO cells in mis-localising MKS3 and CEP290 (Fig. S5g-i). Strikingly, the analysis of NIH3T3 SEPT2 or SEPT7-depleted cells revealed no defect in MKS3 or CEP290 localisation at the TZ (Fig. S5j-k). Together, these data suggest that Septins are required for proper TZ localisation of MKS3 and CEP290 in human but not in mouse cells.

**Figure 5.**
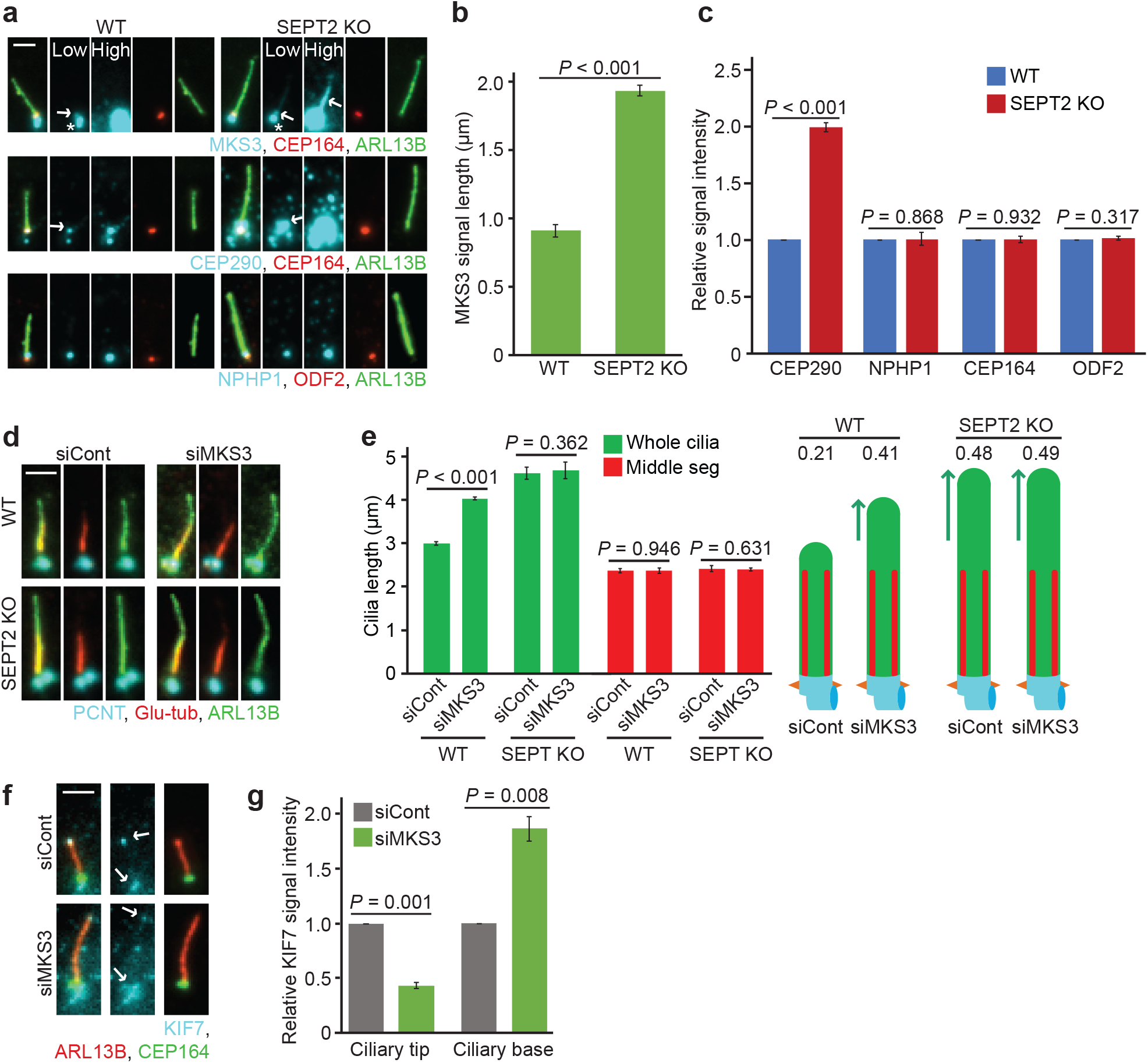
Functional transition zone (TZ) allows to transfer KIF7 to the ciliary tip in RPE1 cells. **a**, Localisation of MKS3, CEP290 and NPHP1 in RPE1 WT and SEPT2-KO after 48h serum starvation (SS) are immunostained with CEP164 (distal appendages), ODF2 (sub-distal appendages), and ARL13B (cilia). White arrows point to the specific signals for MKS3 or CEP290. The white star points to the unspecific MKS3 signal around the centrosome. Low and high exposure times for MKS3m CEP290 and NPHP1 are shown as indicated in the upper panel. Scale bar: 1µm. **b-c**, Quantification of (**a**). The length obtained for MKS3 along the cilia (**b**) and the relative signal intensity of the indicated proteins at the ciliary base (**c**) are shown. Three biological replicates, n=100 cilia per sample and repetition. **d-e**, Influence of MKS3 depletion on cilia length of RPE WT and SEPT2-KO cells. Cells are treated with the indicated siRNA and serum starved (SS) for 48h before immunostaining with the indicated antibodies. Representative images (**d**) and quantifications of middle segment (MS, Glu tub) and whole cilia (ARL13B) length (**e**) are shown. The numbers above the cartoons in (**f**) show the ratio of the calculated DS/whole cilia length. Scale bar: 2 µm. Three 3 biological replicates, n=100 cilia per sample and repetition. **f-g**, A functional TZ is required for KIF7 cilia tip localisation in RPE1 cells. KIF7 signal intensities are measured in ciliated RPE1 cells treated with control or MKS3 siRNA and 48h SS. Representative images (**f**) and quantification of KIF7 signal intensity at the base and tip of cilia (**g**) are shown. Scale bar: 2µm. Three biological replicates, at least 60 cilia per sample and repetition. Data shown in (**b**, **c, e**, **g**) include mean ±s.d. and *P* values are calculated by two-tailed unpaired student t-test. Source data: Supplementary Table 1.

### Depletion of MKS3 causes DS over-elongation and loss of KIF7 at the ciliary tip

We reasoned that accumulation of higher levels of CEP290 at the TZ and MKS3 mis-localisation along the cilia could lead to a non-functional TZ in RPE1 SEPT2-KO cells. In this case, we would expect TZ impairment by MKS3 depletion to phenocopy SEPT2 deletion but not exacerbate the phenotype in SEPT2-KO cells. MKS3-depleted RPE1 cells dropped MKS3 signals at the TZ without decreasing the ciliation rate (Fig. S5a-b, S5l). MKS3-KD significantly extended only the DS in WT but not SEPT2-KO cells (Fig. 5d-e), implying that a functional TZ might contribute to DS hyper-elongation observed in SEPT2-KO cells. In agreement with this assumption, MKS3-KD diminished KIF7 levels at the ciliary tip and increased KIF7 levels at the ciliary base in RPE1 cells (Fig. 5f-g).

We next tested whether MKS3 depletion influences KIF7 localisation in NIH3T3 cells. MKS3-KD in NIH3T3 cells was efficient and did not influence ciliogenesis compared to control cells (Fig. S5m-o). MKS3 depletion led to comparable DS extension in WT and SEPT2-KD cells (Fig. S5p-q). KIF7 at the ciliary tip was markedly decreased in cells lacking MKS3 (Fig. S5r-s).

Together, our data strongly suggest that Septins and a functional TZ are required for proper KIF7 cilia localisation and control of DS length.

### Overgrown cilia dynamically change their length and lose stability in the absence of Septins

To gain insight into DS behaviour, we generated RPE1-WT and SEPT2-KO cells co-expressing ARL13B-GFP and γ-tubulin-mRuby2 to label the cilium and basal body, respectively. Cells were first allowed to ciliate before being subjected to live-cell imaging (Fig. 6a, supplementary movies 1-2). WT cilia maintained an average length around 3µm and the speed of cilia growth and shortening was nearly balanced (Fig. 6a-b, S6a). In contrast, SEPT2-KO cilia showed high length variations (Fig. 6a-b and S6a), which returned back to WT dynamics upon EGFP-SEPT2 expression (Fig. S6b-d, supplementary movies 3-6). During the time of inspection, we also observed two other types of cilia behaviour. In a small percentage of cells, a small vesicle was released from the cilia tip through a process known as ectocytosis (Nager *et al*, 2017; Wood *et al*, 2013) (Fig. 6c, Supplementary movie 5). This event was equally frequent in WT and SEPT2-KO cells (Fig. 6d). In cells with longer cilia, we observed breakage of larger cilia fragments. We refer to these events as excision, as they differed from ectocytosis in respect to the size of released fragments (Fig. 6c-d and Supplementary movie 8). The percentage of excision events was significantly increased in SEPT2-KO compared to WT cells (Fig. 6d) and complementation analysis confirmed that they were related to EGFP-SEPT2 (Fig. S6e). Fluctuations in cilia length and high incidence of excision events were also observed when using SEPT2-KO cells stably expressing SSTR3-GFP as an alternative cilia marker (Fig. S6f-i and Supplementary movies 9-10), indicating that the observed phenotypes were not a consequence of ARL13B-GFP expression.

**Figure 6.**
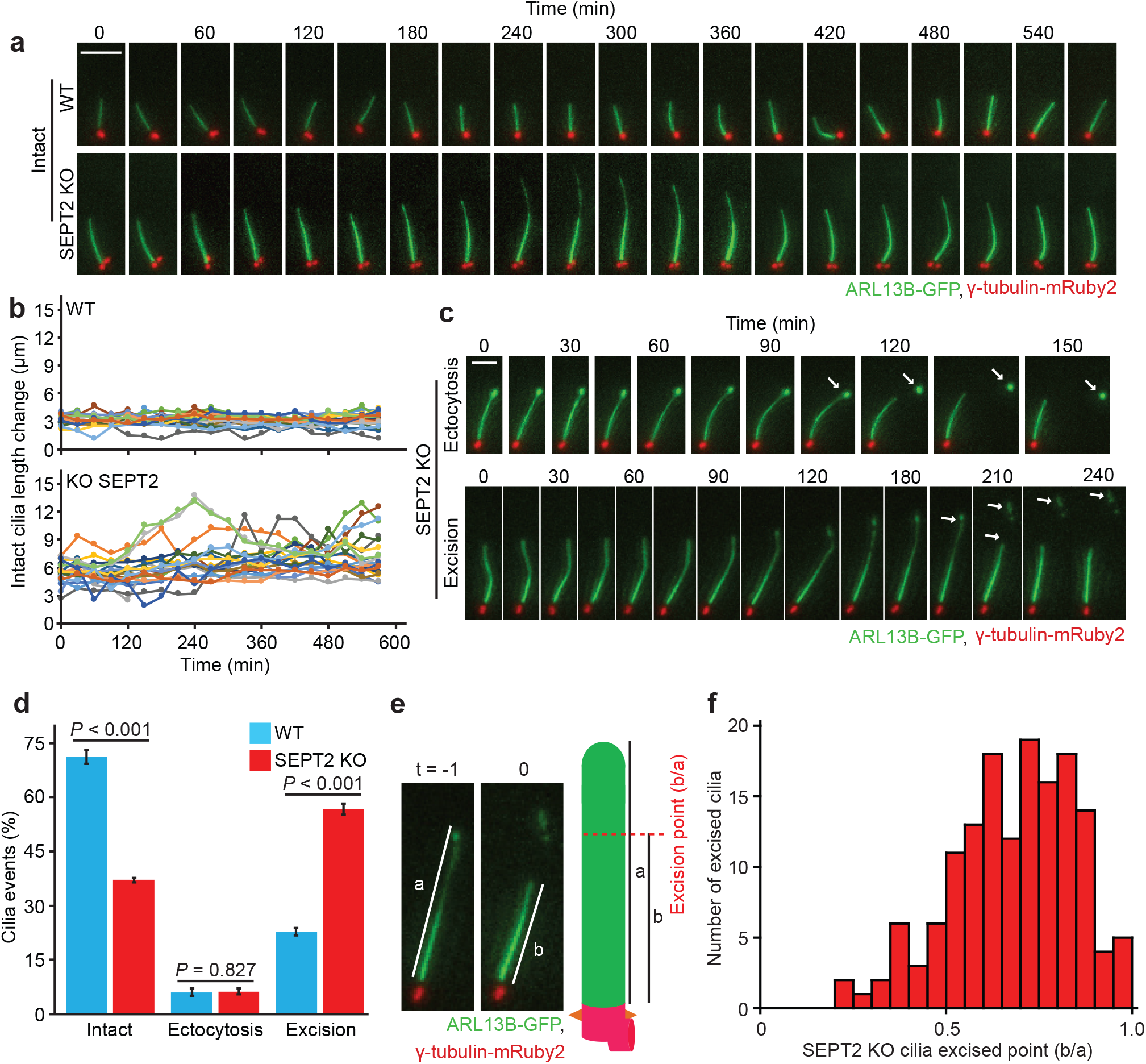
Over-extended distal segments are more prone to undergo excision. **a**, Time-lapse images of RPE1 WT and SEPT2-KO cells stably expressing ARL13B-GFP and γ-tubulin-mRuby2. Cells are starved for 32h before the beginning of inspection and imaged every 30min for 9.5h. Scale bar: 5µm. For videos, see **supplementary movies 1 and 2**. **b**, Quantification of (**a**) showing changes in cilia length during time-lapse imaging. Three biological replicates, 10 cilia per sample and repetition. **c,** Time-lapse images show ectocytosis and cilia excision events observed in RPE1 SEPT2-KO cells. Experiments are done as in (**a**) with exception that cells are imaged every 15min. Scale bar: 2.5µm. For videos, see **supplementary movies 3 and 4**. **d**, Percentage of cilia that remained intact, underwent ectocytosis or excision during live-cell imaging from (**a**) and (**c**). Three 3 biological replicates, n>100 cilia per sample and repetition. **e-f**, Determination of the cilia excision point during live-cell imaging of RPE1 SEPT2-KO ciliated cells. The excision point is calculated by dividing the cilia length at the time of breakage (b, t0) by the length of the cilium one time-point prior to t0 (a, t-1), as depicted in (**e**). The graph in (**f**) shows that breaking points (b/a) higher than 0.5 (i.e. breakage towards the cilia tip) are more frequent. Three biological replicates, n=50 cilia per sample and repetition. Data show in (**d**) mean ±s.d. and *P* values are calculated by two-tailed unpaired student t-test. Source data: Supplementary Table 1.

Next, we analysed the cilia excision phenotype in detail. We hypothesized that the higher rate of cilia excision in SEPT2-KO cells could be linked to DS over-extension. Previous studies reported that the DS is structurally weaker than the MS due to its lower number of axoMTs and fewer structure-strengthening tubulin modifications (Sun *et al*, 2019; Gadadhar *et al*, 2017; He *et al*, 2020). To determine the breakage point, we made use of live-cell images to calculate the excision point (t0) in respect to the total cilia length shortly before breakage (t-1) (Fig. 6e). The quantifications show that excisions happened closer to the cilia distal end (Fig. 6f), indicating that increased excision events in SEPT2-KO cells might be a consequence of a higher number of cilia with over-extended DS. In agreement with this conclusion, we also observed a significant increase in excision events when DS over-extension was induced by nocodazole treatment or upon MAP4, KIF7 or MKS3 depletion (Fig. S6j-k). No increase in excision events was obtained upon CytoD treatment or upon depletion of KIF17 or CCP5, in which the DS was either decreased (KIF17 depletion) or the 30% ratio was maintained (CCP5-depletion, see Fig. 2) (Fig. S6j-k).

Together, our data indicate that Septins are required for cilia length maintenance and prevention of DS instability. It also highlights the importance of keeping a proper DS ratio for cilia stability.

### Balancing DS length is crucial for effective SHH activation

SHH is an established signalling pathway that requires primary cilia for signal transduction in mammalian cells (Ishikawa & Marshall, 2011; Nachury, 2014) (Fig. 7a). We therefore asked whether Septins and DS ratio perturbators, including KIF7 and MKS3, influenced SHH signalling. These studies were done in NIH3T3 cells, which promptly respond to SHH pathway activation after addition of recombinant SHH-ligand (Guo *et al*, 2018; He *et al*, 2014). One of the first events that occurs after binding of SHH-ligands to the receptor Patched-1 (Ptch) is the translocation of the SHH activator Smoothened (Smo) to the cilium (Pedersen *et al*, 2016; Nachury, 2014). In control-depleted cells, cilia accumulation of Smo started shortly after SHH-ligand stimulation and reached a maximum after 4h (Fig. 7b-c). In SEPT2 and SEPT7-depleted cells, the number of Smo-positive cilia was higher compared to control cells at early time points (1-2h, Fig. 7b-c). Percentages of Smo-positive cilia were also increased in MKS3 but not KIF7-KD cells after 2h of SHH-ligand stimulation (Fig. S7a-b). Thus, the initial process of SHH transduction is effective in cells bearing cilia with longer DS.

**Figure 7.**
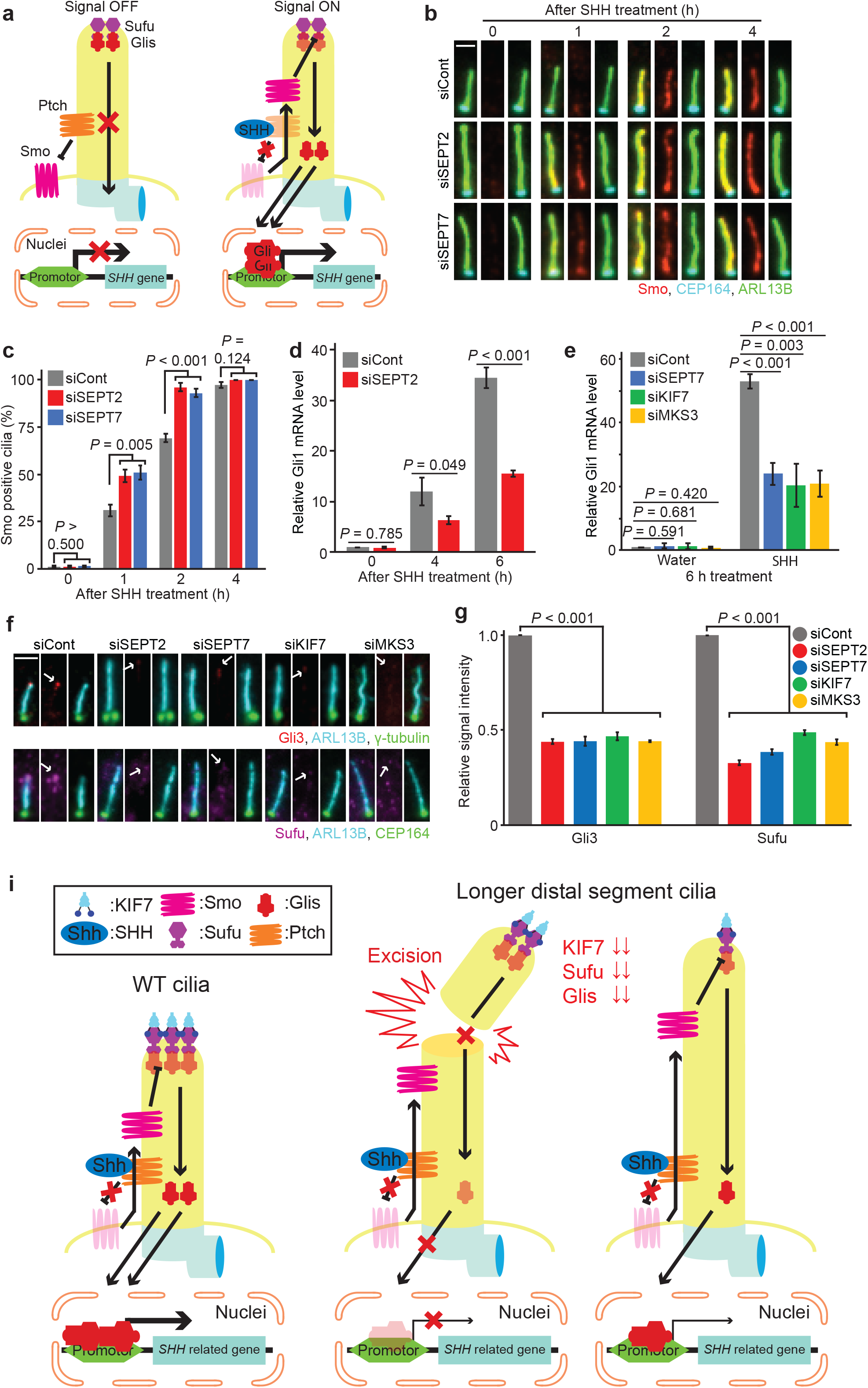
Balancing distal segment length is vital for sonic hedgehog signal transduction. **a**, Schematic representation showing the localisation of SHH components at the cilium in the OFF and ON state(Wheway *et al*, 2018). See text for details. **b-c**, Smo translocation to the cilium changes in the presence or absence of Septins. NIH3T3 cells with control, SEPT2 or SEPT7 siRNAs and 48h serum starvation (SS), and then treated with SHH ligands for 0, 1, 2 and 4h. Smo translocation into the cilium is followed over time using the indicated antibodies. Representative images (**b**) and quantifications (**c**) are shown. Scale bar: 2µm. Three biological replicates, n>100 cilia per sample and repetition. **d**, Relative Gli1 mRNA level changes in control and SEPT2 siRNA-treated NIH3T3 cells. Changes in mRNA level are determined by RT-PCR after 0, 4 and 6h of SHH ligand addition to ciliated NIH3T3 cells. Three biological replicates. **e**, Relative Gli1 mRNA level changes in indicated siRNA-treated NIH3T3 cells. Changes in mRNA level are determined by RT-PCR after 6h of SHH ligand or water addition to ciliated NIH3T3 cells Three biological replicates. **f-g**, Effect of Septins, KIF7 or MKS3 depletion on the localisation of Gli3 and Sufu at the cilia tip. NIH3T3 cells are treated with the indicated siRNAs and 48h SS. Representative images (**f**) and quantifications of Gli3 and Sufu at the cilia tip (**g**) are shown. White arrows in (**f**) point to the specific signal of the indicated proteins at the cilia tip. Scale bar: 2µm. Three biological replicates, n=100 cilia per sample and repetition. **i,** Model depicting the role of longer distal segments on cilia stability and signalling. See text for details. Data shown in **(c, d, e, g)** include mean ± s.d. and *P* values are calculated by two-tailed unpaired student t-test. Source data: Supplementary Table 1.

We asked whether SHH activation downstream of Smo was also faster in SEPT2 absence. For this, we analysed Gli1 transcription factor mRNA levels using real-time quantitative (RT)-qPCR. It is well established that Gli1 mRNA levels rapidly increase upon SHH activation (Shi *et al*, 2015; Taylor *et al*, 2015; Haycraft *et al*, 2005). An increase in Gli1 mRNA levels was observed for control and SEPT2-depleted cells after SHH-ligand stimulation in comparison to non-stimulated cells (t0) (Fig. 7d). However, Gli1 mRNA levels were significantly lower in cells lacking SEPT2 compared to the control (Fig. 7d). Similar results were obtained for SEPT7, MKS3 and KIF7 depletion (Fig. 7e), suggesting that SHH activation is less effective without these components.

To understand the reason for weaker SHH transcriptional activation, we checked the intermediate SHH components: the suppressor of fused (Sufu) and transcription factor Gli3. It is known that Sufu interacts with Gli at the ciliary tip to inhibit Gli translocation into nuclei (Cherry *et al*, 2013). SHH activation relieves this inhibition thereby allowing transcriptional activation (Fig. 7a) (Haycraft *et al*, 2005; He *et al*, 2014; Pedersen *et al*, 2016; Nachury, 2014; Cherry *et al*, 2013). We thus measured the levels of Sufu and Gli3 at the ciliary tip in cells lacking SEPT2, SEPT7, KIF7 and MKS3 (Fig. 7f). In comparison to control cells, Gli3 and Sufu levels dramatically decreased at the ciliary tip in all conditions (Fig. 7f-g). Interestingly, similar results were obtained in cells treated with nocodazole but not CytoD (Fig. S7c-d), implying that like the other components tested, MT perturbation interferes with Sufu and Gli3 localisation at the ciliary tip.

Therefore, we suggest that balancing DS length ratio is required for Sufu and Gli3 cilia localisation and effective SHH activation.

## Discussion

The primary cilium has emerged as an important organelle that controls signal transduction during development and tissue homeostasis. Both, short or abnormally elongated cilia have been reported in patients with ciliopathies and contribute to dysfunctional signalling, highlighting the importance of understanding cilia length regulation (Kim *et al*, 2018; Oh & Katsanis, 2013; Ramsbottom *et al*, 2018; Baala *et al*, 2007; Aguilar *et al*, 2012). Whereas the majority of studies were so far dedicated to the analysis of cilia assembly and disassembly, little is known about how cilia length is maintained at steady-state. Here, we characterized cilia behaviour based on different cilia markers and established Septins as important regulators of cilia length, in particular in the stability of the DS in human and mouse cells.

Analysis of cilia segmentation based on cilia membrane localisation and tubulin glutamylation indicated that RPE1 and NIH3T3 ciliated cells contain polyglutamylated-rich and -poor regions, referred to as the middle and distal segments respectively, in analogy to previous work done in *C. elegans* and IMCD3 cells (Kramer *et al*, 2007; Sun *et al*, 2019). Our data suggest that the DS is formed after MS elongation, implying a time-dependent maturation of the cilium. At steady-state, the DS of RPE1 and NIH3T3 cilia reached approximately 30% of the entire cilia length, whilst the remaining 70% were formed by the MS. Intriguingly, treatments leading to cilia over-elongation disturbed this ratio in different ways. For instance, low doses of nocodazole or depletion of cilia proteins DYNC2L1, KIF3A, MAP4, SEPT2 or MKS3 led to DS elongation without affecting MS length. As a consequence, the DS:MS ratio was shifted to 50:50 under these conditions. On the other hand, actin depolymerisation by CytoD or CCP5 depletion also increased MS length, however the DS:MS ratio remained at 30:70. Although the cilium was overall longer under all these conditions, the main difference was a change in the DS:MS ratio. Our data suggest that keeping a balanced DS:MS ratio might be key for determining cilia stability and behaviour. In all conditions analysed here, in which the DS exceeded 30% of cilia length, we observed an increase in excision events releasing larger cilia fragments. We did not observe increased excision in CytoD treated and CCP5-KD cells, in which the cilium was hyper-elongated, yet the DS:MS ratio of 30:70 was preserved. This indicates that cilia have a built-in ruler that measures and adjusts the relative length of ciliary subdomains to maintain its stability.

Septins were reported as positive regulators of cilia formation acting at the ciliary base in IMCD3 cells or along the axoneme in RPE1 cells. In both cases, Septins were proposed to control ciliary composition and MT behaviour by either functioning as a diffusion barrier at the base of the cilium or competing with MAP4 for axoneme binding. Our study revises these models and now reveals that Septins are involved in controlling DS stability. Several lines of evidence support this conclusion. The depletion of multiple Septins in RPE1 and NIH3T3 cells led to cilia DS over-extension. This phenotype was recapitulated using distinct RPE1 SEPT2-KO cell lines. In all cases, cilia over-extension was rescued by exogenous Septin-carrying constructs, implying that the effect on cilia length control was specific to Septins. Moreover, electron microscopy confirmed the presence of hyper-elongated cilia in SEPT2-KO cells. Live-cell imaging revealed an increased rate of cilia elongation and of excision of larger DS parts, leading to rapid cilia shortening. These data contrast from the previous conclusion that Septin depletion generally leads to assembly of shorter cilia. One possibility is that Septins might play different roles depending on cell type. Alternatively, as previous studies mainly used end-point analysis of fixed samples, we envisage that experimental conditions during sample handling could exacerbate cilia breakage in Septin-less cells, masking the initial DS hyper-elongation phenotype. Nevertheless, these data agree with the fact that Septins play an important role in cilia maintenance by keeping the proper cilia length at steady-state.

Our data support the model that Septins restrict DS elongation by promoting localisation of KIF7 (Fig. 7i). KIF7 was shown to bind to the plus-end of MTs at the ciliary tip, forming a cap structure that restricts axoMTs extension (He *et al*, 2014). Loss of KIF7 at the ciliary tip in cells lacking Septins most likely accounts for DS de-regulation. In human cells, KIF7 accumulated at the ciliary base, implying a TZ defect. The mis-localisation of MKS3, CEP290 (this study) and TMEM231 (Chih *et al*, 2012) in RPE1 cells argues in favour of our hypothesis. Furthermore, MKS3-KD in RPE1 and NIH3T3 cells caused DS over-elongation and KIF7 loss from the ciliary tip, emphasizing that a functional TZ is essential for proper control of KIF7 localisation and cilia growth.

How do Septins contribute to TZ assembly and KIF7 localisation? Septins localise along the cilia in RPE1 cells, but not equally. The co-localisation analysis of SEPT2 and TZ components showed that SEPT2 levels were lower at the TZ in comparison to other segments. One possibility is that Septins create a membrane diffusion barrier at the cilia to confine TZ components in place. This function would be in line with the fact that membrane-bound TZ components, MKS3, CEP290 and TMEM231, failed to be retained at the ciliary base and extended along the cilia (this study and (Chih *et al*, 2012)). However, as mis-localisation of KIF7 but not TZ components was observed in NIH3T3 cells, it is possible that Septins contribute to KIF7 cilia localisation independently of its function in TZ organisation. Considering that Septins are widespread in the cytoplasm of RPE1 and NIH3T3 cells, it is also feasible that Septins fulfil functions outside the cilium that could influence protein transport to the basal body.

By analysing components of the SHH signalling pathway, we observed different behaviour of SHH components in response to longer-distal segment forming cells. Shortly after SHH activation, SMO entered the cilium in both control and Septins-, KIF7-, and MKS3-depleted conditions, showing responsiveness to SHH activation. However, the Gli and Sufu levels at the ciliary tip were drastically decreased in the absence of those genes, explaining the impairment in the downstream activation of SHH, when using Gli1 mRNA levels as a read-out. KIF7 physically interacts with Glis, Sufu and Smo to modulate their function in signalling transduction (Chih *et al*, 2012; Cheung *et al*, 2009; Li *et al*, 2012; Endoh-Yamagami *et al*, 2009). The lack of KIF7 at the cilium in cells lacking Septins could explain the reduction in the downstream SHH signalling response.

Analysis of cilia derived from patients carrying ciliopathies, such as Meckel and Joubert syndrome, show that inactivating mutations of MKS3 and CEP290, respectively, form abnormally elongated cilia and severe defects in Shh pathway activity (Aguilar *et al*, 2012; Ramsbottom *et al*, 2018; Tammachote *et al*, 2009; Srivastava *et al*, 2017). Our live-cell images indicate that cells with abnormally elongated cilia cannot stabilize the cilia, leading to increased distal breakage, where most of the signalling receptors accumulate (Wheway *et al*, 2018; Nager *et al*, 2017; Haycraft *et al*, 2005). Therefore, it is tempting to speculate that abnormally long cilia might impair signalling through increased excision events and loss of signalling receptors from the cilium.

Cilia length was shown to vary depending on the cell type (Besschetnova *et al*, 2010; Miyoshi *et al*, 2014; Qiu *et al*, 2012; Wann & Knight, 2012). It will be interesting to determine whether the 30:70 DS:MS ratio also applies to various types of primary cilia and whether mutations that perturb this ratio lead to cilia instability and signalling activation impairment, similar to the phenotypes described here for human and murine epithelial cells.

## Materials and methods

### Plasmids and Reagents

Plasmids used in this study are listed in Supplemental table S2. Human SEPT2 (BC014455) and SEPT6 (BC009291) cDNA were obtained from DKFZ (Clone 121547190 and 121516346, respectively). Human SEPT7 (BC093642.1) was purchased from Dharmacon (Cat. MHS6278-211688440). For siRNA rescue experiments in RPE1 cells, siRNA-resistant SEPT2, SEPT6 and SEPT7 GFP-fusion constructs were generated by introducing 3 silent point-mutations at the siRNA targeting sites by site-directed PCR mutagenesis (Table S2). All reagents and media were purchased from Sigma-Aldrich unless specified otherwise. Recombinant Human SHH Protein for SHH pathway activation was purchased from R&D systems.

### Cell culture and treatments

Tet3G integrated human telomerase (hTERT)-immortalised retinal pigment epithelial cells (RPE1-Tet3G) (Hata *et al*, 2019) (kind gift from Elmar Schiebel, University of Heidelberg, Germany) were cultured in DMEM/F12 supplemented with 10% fetal bovine serum (FBS, Biochrom), 2 mM L-glutamine (Thermo Fischer Scientific) and 0.348% sodium bicarbonate. Mouse immortalized fibroblast NIH3T3 cells (ATCC, CRL-1658) were grown in DMEM high glucose supplemented with 10% new-born calf serum (NCS, PAN-Biotech). HEK293T (ATCC CRL-3216) and GP2-293 cells (Takara Bio, Cat. 631458) were maintained in DMEM high glucose supplemented with 10% FBS. All cell lines were grown at 37°C with 5% CO_2_. siRNA-based gene knockdown was performed with Lipofectamine RNAiMAX transfection reagent (Thermo Fischer Scientific) and 20 nM (final concentration) of corresponding siRNAs accordingly to the manufacture protocol. The list of siRNAs used in this study can be found in Supplemental Table S3.

For induction of ciliogenesis, cells were seeded in 24-well plates (30.000 cells/well) in serum-rich media. Twenty-four hours after seeding, the medium was exchanged to medium lacking FBS (RPE1) or containing 0.5 % NCS (NIH3T3). Cultures were inspected up to 48h after serum starvation. For cilia analysis after gene knockdown, treatment of cells with siRNA was done 24h prior to serum starvation. For chemical treatments, cells were starved for 48h following treatment with DMSO (solvent control), Cytochalasin D (final concentration: 200 nM) and/or nocodazole (final concentration: 100 nM) for 3h before inspection. For Shh pathway activation, NIH3T3 cells were starved for 48h and treated with 50 pM Shh ligand (final concentration) for the indicated time points.

### Generation of stable cell lines

Stable RPE1-Tet3G cell lines were generated by virus-based gene integration. HEK293T or GP2-293 cells were used for lentivirus or retrovirus production using polyethyleneimine (PEI 25000, Polysciences). Supernatants containing lentiviral and/or retroviral particles were applied to RPE1-Tet3G-derived host cells for 24h. The following cell lines were generated: RPE1-Tet3G constitutively expressing ARL13B-GFP and γ-Tubulin-mRuby2; RPE1-Tet3G constitutively expressing SSTR3-GFP and γ-Tubulin-mRuby2; RPE1-Tet3G constitutively expressing ARL13B-mRuby2; RPE1-Tet3G expressing siRNA-resistant EGFP-SEPT2, EGFP-SEPT6 or SEPT7-EGFP under control of the inducible Tet-promoter; RPE1-Tet3G ARL13B-mRuby2 with siRNA-resistant EGFP-SEPT2 under control of the inducible Tet-promoter. Low levels of fluorescence positive cells were sorted by fluorescence activated cell sorting (FACS) (BD FACS Aria III, Becton Dickinson). For cell lines expressing constructs under the Tet-promoter, cells were sorted for low GFP expression levels 24h after treating the cells with 10 ng/ml doxycycline.

### Generation of knockout cell lines by CRISPR/Cas9 system

CRISPR/Cas9-mediated chromosomal deletion was used for generating knockout human SEPT2 cells in RPE1 Tet3G cells. Two gRNAs targeting human SEPT2 exon 4 (5’-ggggttcgagttcactctgatgg-3’) and 6 (5’-ccggctacggggatgccatcaac-3’) were cloned into Cas9 system expression plasmid pX458 to generate pTK58 and pTK60 (Table S2). RPE1 Tet3G cells were transfected with pTK58 or pTK60 by electroporation (Neon® Transfection System) according to the manufacture’s protocol. One day after electroporation, GFP positive cells were FACS sorted and subjected to single cell dilutions. Single clones were allowed to expand for 10-14 days before they were subjected to immunoblotting to check SEPT2 protein levels as well as genomic amplification followed by cloning into pJET vector (Thermo Fischer Scientific). The plasmids were then sequenced to identify clones containing two SEPT2 mutated alleles with premature stop codons (Individual clone information is shown in Fig. S3f). Two independent clones were selected per gRNA for further analysis.

### Antibodies

Primary antibodies used in this study can be found in Table S4. Secondary antibodies used for wide-field microscopy were purchased from Thermo-Fisher Scientific and included: goat anti-rabbit, anti-mouse or anti-guinea pig conjugated to Alexa Fluor 350, 488, 594 or 647 (1:500 dilution); donkey anti-rabbit, anti-mouse or anti-goat conjugated to Alexa 488, 555 and 594 (1:500 final dilution). For STED microscopy, anti-rabbit Atto594 (1:100 dilution) and anti-mouse Star635p (1:100 dilution) were purchased from Sigma and Abberior, respectively. Secondary antibodies for immunoblotting were horseradish peroxidase conjugated goat anti-rabbit or anti-mouse antibodies (1:1000 dilution) (purchased from Jackson ImmunoResearch).

### Immunofluorescence microscopy (IF)

Cells were cultured on coverslips (No. 1.5, Thermo Fischer Scientific) and fixed for IF as following: cells carrying EGFP-tagged Septins and ARL13B-mRuby2 were fixed in 3% paraformaldehyde at room temperature for 5 min and cold methanol at −20°C for 5 min; in all other experiments, cells were only fixed in cold methanol at −20°C for 5 min. After fixation, cells were treated with phosphate buffer (PBS) containing 3% BSA (Jackson Immuno Research) and 0.1% Triton X-100 (blocking solution) for 30 min at room temperature in a wet-chamber. Blocked samples were incubated with primary antibodies diluted for 1h at room temperature. After washing with PBS, samples were incubated with secondary antibodies for 30 min at room temperature. For DNA stainings, DAPI (4’, 6-diamidino-2-phenylindole) at a final concentration of 1 µg/ml was added to the secondary antibody dilution. All antibodies were diluted in blocking solution. Coverslips were mounted with Mowiol (EMD Millipore).

Wide-field fluorescence microscopy images were acquired as Z-stacks using these microscopes; Nikon Eclipse Ti2 Inverted Microscope Systems with Plan Apo 40x/0.95 and 60x/0.45 Oil objectives and an IRIS9 Scientific CMOS camera operating Nikon NIS-Elements Imaging Software; Zeiss Axiophot with 63× NA 1.4 Plan-Fluor oil immersion objective, and Cascade:1K EMCCD camera operating Meta.Morph software; Zeiss Axio Observer Z1 with 63× NA 1.4 Plan-Apochromat oil immersion objective, and AxioCam MRm CCD camera operating ZEN software. For cilia length measurements and the number of ciliated cells, quantifications were done using Z-stacked max-signal intensity projected images. Relative fluorescence intensities of targeted proteins were quantified using Fiji ImageJ software (Schindelin *et al*, 2012). Signal intensities were normalized by dividing to control average intensities of each experiment.

STED images were acquired on Leica TCS SP8 STED 3X super-resolution microscope platform mounted on Leica DMi8, inverted microscope with HC PL APO 100x/1.40 STED White Oil objective, 2 Hybrid GaAsp detectors (HyD) and 2 PMTs. The imaging software was controlled by Leica Application Suite 3 (LAS) and image processing was performed by Huygens deconvolution pro and Fiji.

Images were adjusted for contrast or brightness in Fiji. Figures were assembled in Adobe Photoshop and Illustrator CS3 (Adobe).

### Electron microscopy

RPE1-Tet3G and RPE1-Tet3G SEPT2-KO cells were serum starved for 48h to induce ciliogenesis. Cells were rinsed with 100 mM PBS 3 times and then fixed with a mixture of 2.5 % glutaraldehyde, 1.0 % PFA and 2 % sucrose in 50 mM cacodylate buffer for 30 min at room temperature. The fixative was washed out with 50 mM cacodylate buffer. After post-fixation with 2 % OsO_4_ for 1h at 4°C in darkness, cells were rinsed 4 times with water and incubated overnight at 4°C in aqueous 0.5 % uranyl acetate solution. On the following day, coverslips were washed 4 times with water and dehydrated with consecutive incubations in ethanol solutions (40, 50, 70, 80, 90, 95, 100 %). Coverslips were immediately placed on EM-capsules filled with Spurr-resin and allowed to polymerize at 60°C for 24-48h. Embedded cells were sectioned using a Reichert Ultracut S Microtome (Leica Instruments, Vienna, Austria) to a thickness of 80 nm. Post-staining was performed with 3 % uranyl acetate and lead citrate. Serial-sections were imaged at a Jeol JE-1400 (Jeol Ltd., Tokyo, Japan), operating at 80 kV, equipped with a 4k x 4k digital camera (F416, TVIPS, Gauting, Germany). Micrographs were adjusted in brightness and contrast using Fiji.

### Immunoblotting

For immunoblot analysis, whole cell lysates were obtained by incubating the cell pellets with 8 M Urea containing benzonase (1:1000 dilution) at room temperature for 1h. The protein concentration was measured using Bradford reagent accordingly to manufactures instructions. Equal amount of protein was loaded on SDS–PAGE. Separated proteins were transferred onto PVDF membranes, then blocked for 30 min in PBS containing 5% milk and 0.1% Tween-20. Membranes were incubated with primary (4°C overnight) and secondary antibodies (1 h, room temperature) dilutions and proteins were visualized with enhanced chemiluminescence (ECL) (Pierce; Thermo Fisher Scientific). For loading control (Supplementary figures 3b and 3k), detection of actin was performed on stripped membranes (Stripping solution: 1% SDS, 0.2M Glycine pH 2.5 in water).

### Live-cell imaging

For live-cell imaging, cells were cultured in HEPES-buffered DMEM/F12 supplemented with 10% FBS, 1% L-glutamine and 1% penicillin–streptomycin without phenol red (ThermoFischer) in glass-bottom CELLview culture dishes (Greiner Bio-One) for 24h and serum starved for 32h. Dishes were mounted in a pre-heated microscope chamber and live cell imaging was performed over a period of 15h at 37°C with 5% CO_2_. 3D-Images (8 z-stacks of 6 µm spacing) were acquired by the Nikon/Andor TuCam system mounted with the new Andor Neo sCMOS camera and a Nikon Plan Apo VC 60x NA 1.4 oil immersion objective every 15 min. For chemical treatment, 200 nM Cytochalasin D or 100 nM nocodazole was added 30 min before imaging. For rescue experiments, 10 nM doxycycline was added to induce EGFP-SEPT2 protein expression when cells were initially seeded on CELLview culture dishes. Images were processed in Nikon NIS-Elements Imaging Software using maximum intensity projections. The cilia length in each time-point was manually measured using Fiji.

### Real-time quantitative PCR (qPCR)

mRNA was isolated using NucleoSpin® RNA Plus (Machery-Nagel) and 1 µg of total mRNA was reverse-transcribed into cDNA using SensiFAST^TM^ cDNA Synthesis Kit (Bioline) according to the manufacture’s protocol. The cDNA (100 ng) was analysed by quantitative PCR using SensiFAST^TM^ SYBR Lo-ROX Kit (Bioline) in the 7500 Fast Real-Time PCR System (Applied Biosystems). PCR products were amplified using following primer sets: mGli1 Fw: 5’-GAGAAGCCACACAAGTGCACGT-3’, mGli1 Rv: 5’-AGGCCTTGCTGCAACCTTCTTG-3’, mGapdh Fw: 5’-CATCACTGCCACCCAGAAGACTG-3’, mGapdh Rv: 5’-ATGCCAGTGAGCTTCCCGTTCAG-3’. qPCR data were analysed using the ΔΔCT methods and normalized against the house-keeping gene GAPDH(Shi *et al*, 2015). The RQ value (fold change) was calculated using the formula 2^(-ΔΔCT)^.

### Statistics and reproducibility

Statistical analyses were done using Wilcoxon or two-tailed Student’s t-tests. Significance probability values were P < 0.05. Statistical tests were performed in Excel (Microsoft) and KaleidaGraph (Synergy Software). Statistical analysis and mean ± standard deviation (sd) were performed on at least three independent biological replicates done in similar conditions. The measurement of cilia length by electron microscopy and data shown in figures S6c and S6d were performed two times independently. The number of biological replicates and sampling sizes are indicated in figure legends and Supplemental Table S1.

## Acknowledgements

We thank E. Schiebel (ZMBH, Heidelberg University, Germany) for sharing cell lines, antibodies and plasmids, K. Anderson (Memorial Sloan-Kettering Cancer Centre, USA) for sharing KIF7 antibody, Astrid Hofmann (member of the Pereira lab) for technical support, H. Lorenz (ZMBH Imaging facility, Heidelberg University, Germany) for helping with STED microscopy, U. Engel (Nikon Imaging Centre, Heidelberg University) for sharing the Nikon/Andor TuCam microscope for live-cell imaging, T. Holstein (COS, Heidelberg University, Germany) for access to the qPCR machine and M. Langlotz (ZMBH FACS facility, Heidelberg University, Germany) for cell sorting. We thank E. Schiebel and H. Lee (member of the Pereira lab) for critically reading the manuscript, all members of the Pereira lab and, especially, S. Hata (University Tokyo, Japan) for helpful discussions. TK received financial support from the Heidelberg Biosciences International Graduate School (HBIGS). Funding was obtained from the collaborative research grant of the Deutsche Forschungsgemeinschaft (DFG) (SFB873, Project A14) granted to GP. Core funding for microscopy and FACS was provided by the SFB873 Projects Z03 and Z02, respectively. The work of GP was supported by the Deutsche Forschungsgemeinschaft Heisenberg Professorship (PE1883/3, PE1883/4). The authors declare no competing financial interests.

## Author contributions

G.P. conceived and supervised the project. T.K. designed, performed the experiments and analysed the data. A.N. performed experiments related to electron microscopy. B.K. supported the experiments. G.P. and T.K. wrote the manuscript. All authors contributed to manuscript preparation and discussion.

## Supplemental figure legends

**Figure S1.**
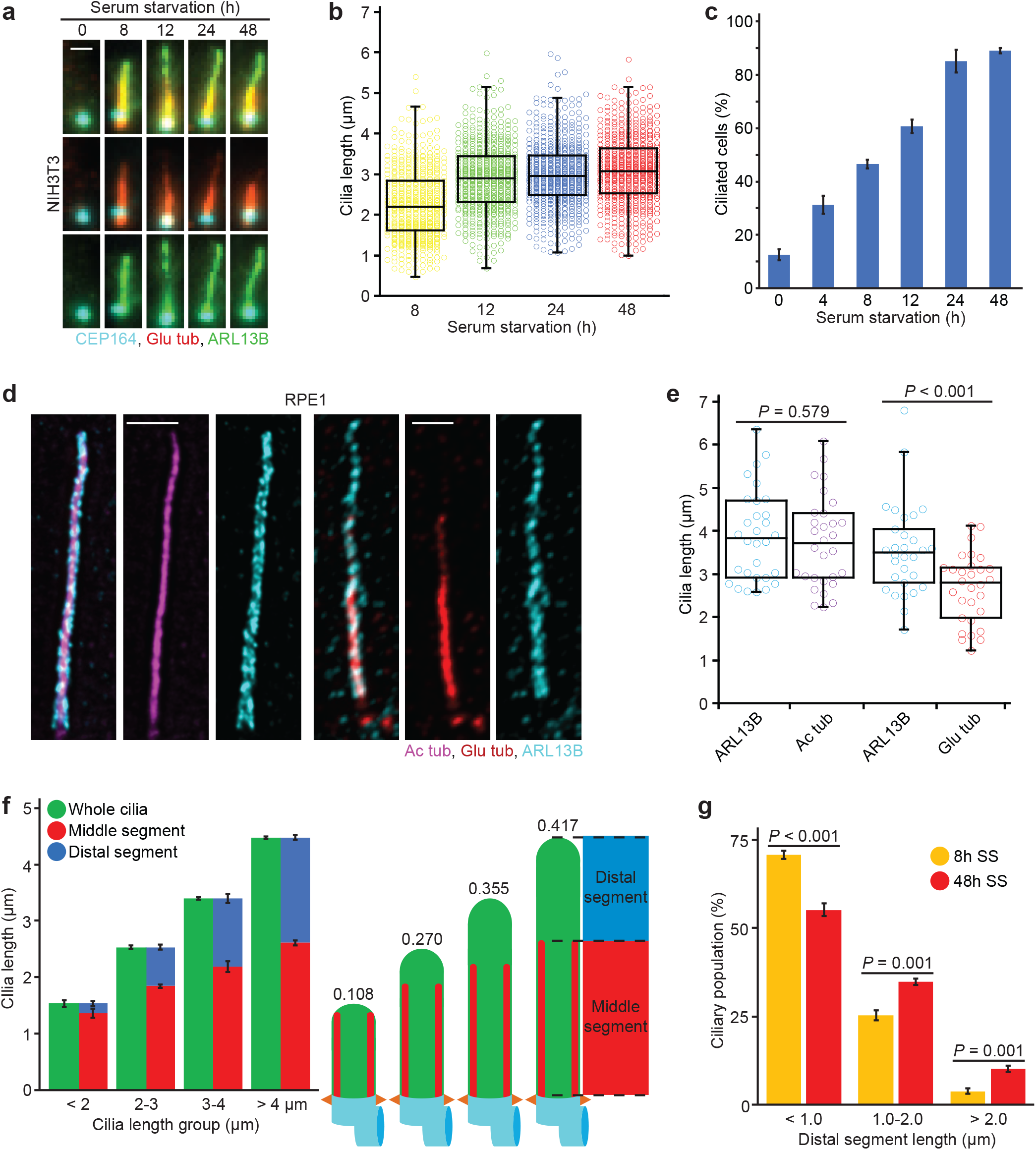
Characterisation of middle/distal cilia segmentation in RPE1 and NIH3T3 cells. **a-b**, Representative images of cilia in NIH3T3 cells (**a**) and dot plots (**b**) showing primary cilia length variation at different time points after serum starvation (SS). Markers in (**a**) show ARL13B (cilia), polyglutamylated tubulin (Glu tub, middle segment) and CEP164 (Basal body). The graph shows individual length of cilia from 3 biological replicates, n>150 cilia per sample and experiment. Scale bar: 1µm. **c**, Percentage of ciliated NIH3T3 cells after SS from **a**. 3 biological replicates, n>100 cells per sample and repetition. **d**, STED microscopy images showing ciliated RPE1 cells stained with acetylated tubulin, Glu tub and ARL13B. Scale bar: 1µm. **e**, Quantification of (**d**) comparing cilia length as determined by the indicated staining. n=30 cilia from three biological replicates. **f**, Average length of MS and DS for the cilia length groups depicted. Representative images (**d**) and quantifications of MS (Glu tub) and whole cilia (ARL13B) length are shown. The numbers above the cartoons show the ratio of the calculated DS/whole cilia length. Three biological replicates, n>700 cilia per sample and repetition. **g**, Percentage of cilia with depicted DS length after 8h and 48h SS. Three biological replicates, n>150 cilia per sample and repetition. Data shown in **(b, c, e, f** and **g)** include mean ±s.d.; *P* values are calculated by unpaired Wilcoxon-Mann-Whitney Rank Sum Test (in **b, e)** or two-tailed unpaired student t-test (in **c, g**). Source data is shown in Supplementary Table 1.

**Figure S2.**
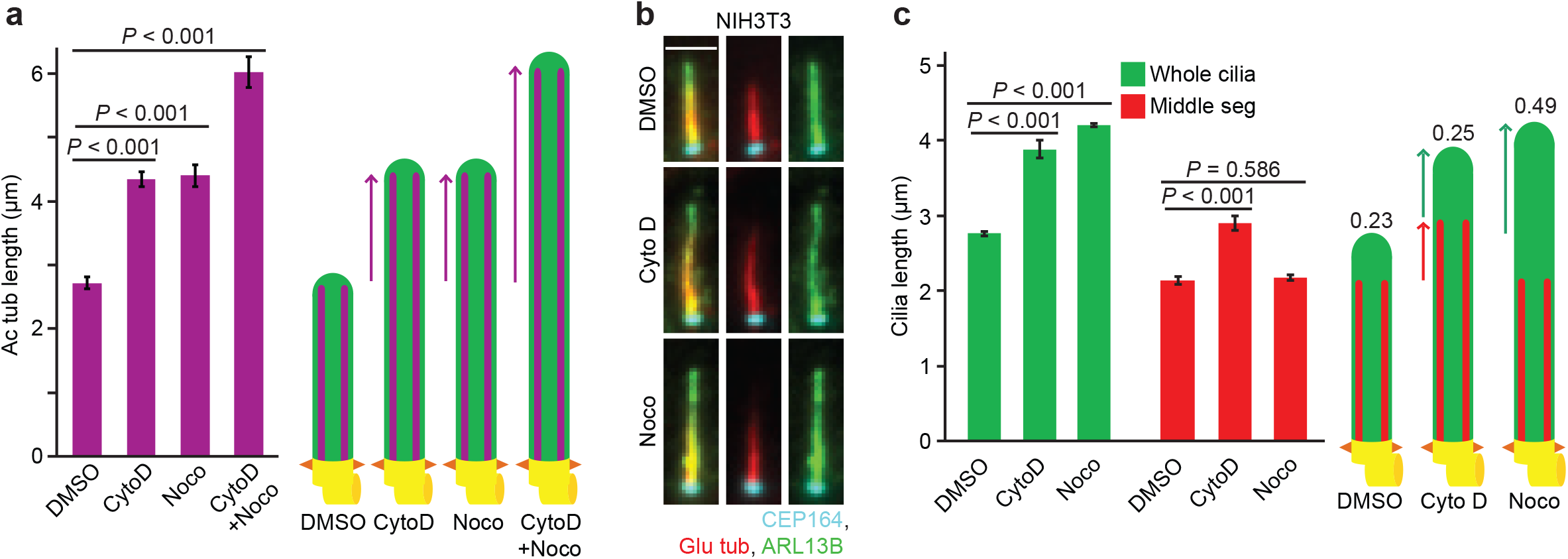
Microtubule and actin perturbators change cilia segmentation in the different ways. **a,** Representative images of 48h serum starved RPE1 cells treated with solvent control (DMSO), cytochalasin D (CytoD) and/or nocodazole (Noco) for 3h. The length of cilia was determined based on acetylated tubulin (Ac tub) staining. Three biological replicates, n=100 cilia per sample and repetition. This data is related to Fig. 2a. **b,** Representative images of 48h serum starved NIH3T3 cells treated with solvent control (DMSO), cytochalasin D (CytoD) and/or nocodazole (Noco) for 3h before immunostaining with the indicated antibodies. Scale bar: 2µm. **c,** Quantification of (**b**) showing middle segment (MS, Glu tub) and whole cilia (ARL13B) average length. The numbers above the cartoons show the ratio of the calculated distal segment (DS)/whole cilia length. Three biological replicates, n=100 cilia per sample and repetition. Data shown in **(a, c)** include mean ±s.d. and *P* values are calculated by two-tailed unpaired student t-test. Source data is shown in Supplementary Table 1.

**Figure S3.**
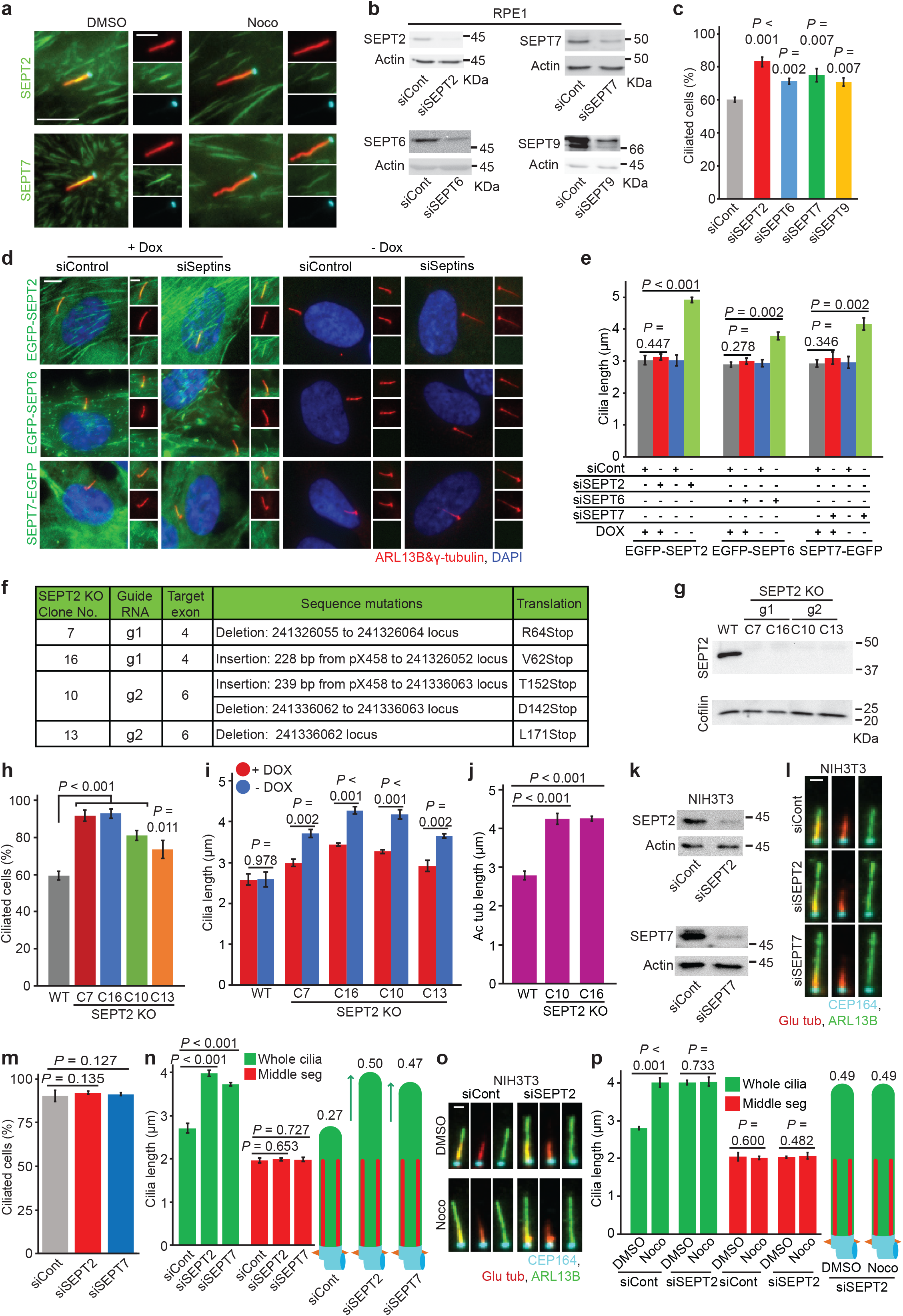
Septin negatively controls distal segment elongation in human and mouse cells. **a,** Effect of nocodazole (Noco) treatment (3h) on the localisation of endogenous SEPT2 and SEPT7 at cilia in RPE1 cells. The insets show magnifications of the cilium. Scale bar: 5µm (large) and 2.5µm (small). **b**, Immunoblot analysis to assess the efficiency of siRNA depletions in RPE1 cells. Anti-SEPT2, SEPT7, SEPT6, SEPT9 and actin antibodies are used. Actin served as a loading control. **c,** Percentage of ciliated RPE1 cells with indicated siRNA-treatment from Fig. 3e. Three biological replicates, n>100 cells per sample and repetition. **d-e**, Expression of siRNA-resistant GFP-tagged Septin constructs rescue cilia length in RPE1 cells. The expression of the constructs is induced by addition of doxycycline (DOX) as indicated. Representative images (**d**) and quantifications (**e**) are shown. The insets in (**d**) show magnifications of the cilium. Scale bar: 5µm (large) and 2.5µm (small). Three biological replicates, n>80 cilia per sample and repetition. **f**, Table summarises the genomic information of RPE1 SEPT2-KO clones. **g**, Immunoblot shows SEPT2 protein levels in RPE1 WT and SEPT2-KO cells. Anti-SEPT2 and Cofilin antibodies are used. Cofilin served as a loading control. **h,** The percentage of ciliated RPE1 WT and SEPT-KO cells from Fig. 3g. Three biological replicates, n>100 cells sample and repetition. **i,** Expression of GFP-SEPT2 after DOX addition reduces cilia length in RPE1 SEPT2-KO cells. Three 3 biological replicates, n>80 cilia per sample and repetition. **j,** Length of acetylated tubulin (Ac tub) in RPE1 WT and SEPT2-KO cells from Fig. 3i. Three biological replicates, n=100 cilia per sample and repetition. **k**, Immunoblot analysis to assess the efficiency of siRNA depletions in NIH3T3 cells. Anti-SEPT2, SEPT7 and actin antibodies are used. Actin served as a loading control. **l-n**, NIH3T3 cells are treated with the indicated siRNAs and serum starved for 48h. Cilia are stained with polyglutamylated tubulin (Glu tub, middle segment), ARL13B (cilia) and CEP164 (basal body). Representative images (**l**), percentage of ciliated cells (**m**) and segmentation analysis (**n**) are shown. The numbers above the cartoons in (**n**) indicate the ratio of the calculated distal segment (DS)/whole cilia length. Scale bar: 1.5µm. Three biological replicates, n>100 or n=100 cells per sample and experiment were quantified in (**m)** or (**n**) respectively. **o-p**, Ciliated NIH3T3 cells with indicated siRNA-treatment are treated with solvent control (DMSO), cytochalasin D (CytoD) and/or nocodazole (Noco) for 3h before immunostaining with the indicated antibodies. Representative images (**o**) and quantifications of MS (Glu tub) and whole cilia (ARL13B) length (**p**) are shown. The numbers above the cartoons in (**p**) show the ratio of the calculated DS/whole cilia length. Scale bar: 2µm. Three biological replicates, n=100 cilia per sample and repetition. Immunoblots shown in **(b, g, k)** are replicated twice. Data shown in **(c, e, h, i, j, m, n** and **p)** include mean ±s.d. and *P* values are calculated by two-tailed unpaired student t-test. Source data is shown in Supplementary Table 1.

**Figure S4.**
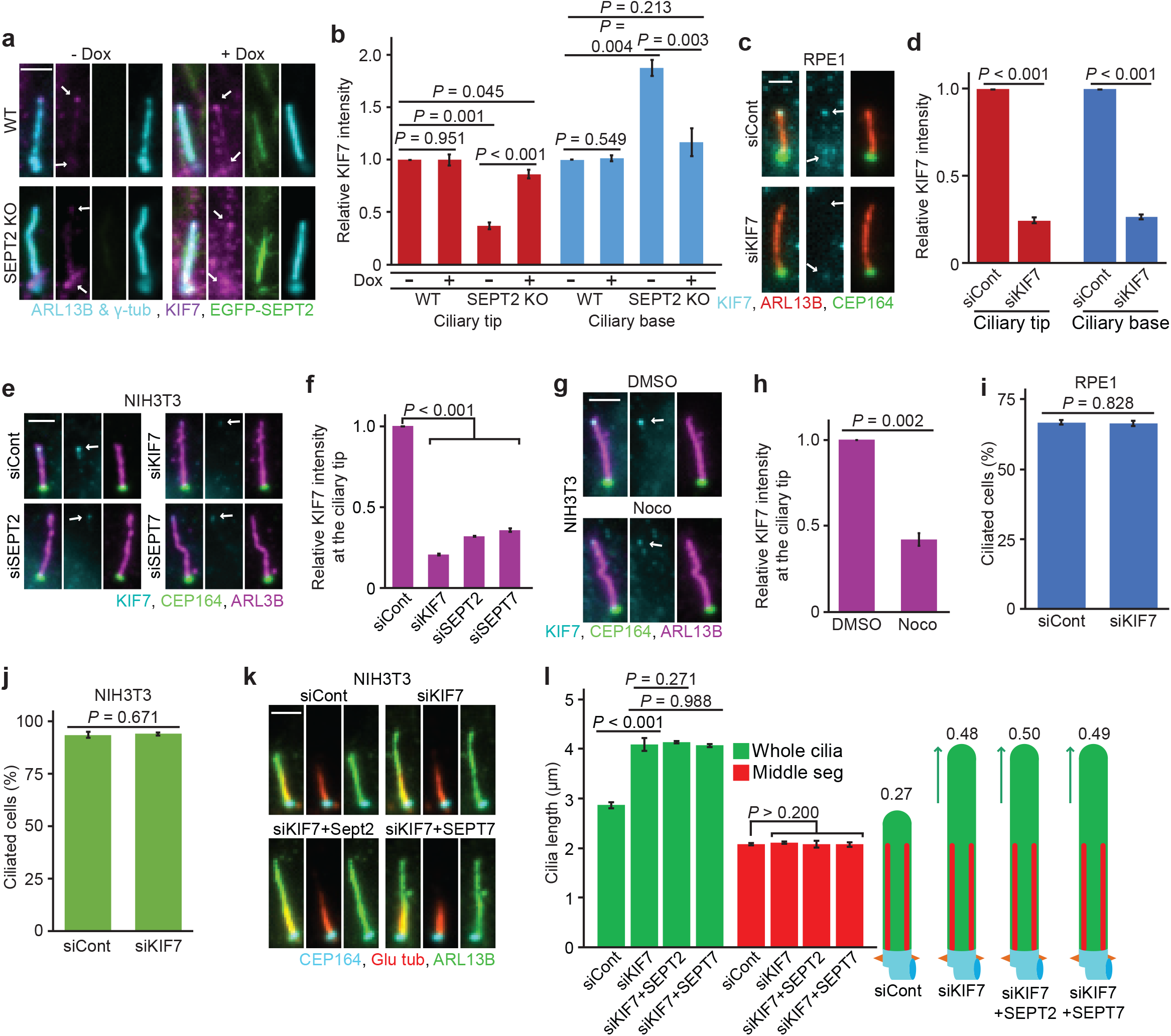
Septin is required for KIF7 accumulation at the cilia tip to suppress distal segment growth. **a-b**, Rescue experiment of KIF7 localisation at the cilia after expression of GFP-SEPT2 in RPE1 SEPT2-KO cells. GFP-SEPT2 expression was induced by addition of doxycycline (DOX) as indicated. Representative cells (**a**) and quantifications of the relative signal intensity of KIF7 at the tip and base of the cilia (**b**) are shown. The arrows in (**a**) point to the ciliary base and tip. Scale bar: 2µm. Three biological replicates, n>80 cilia per sample and repetition. **c-d**, Efficiency of KIF7 knockdown in RPE1 cells. Representative images (**a**) and quantifications of KIF7 protein levels at the tip and base of cilia (**b**) are shown. The arrows in (**c**) point to the cilia base and tip. Scale bar: 2µm. Three biological replicates, at least 60 cilia per sample and experiment. **e-f**, KIF7 localisation changes in NIH3T3 cells treated with control, KIF7, SEPT2 or SEPT7 siRNAs. Representative images (**e**) and quantifications of KIF7 signal intensities at the cilia tip (**f**) are shown. The arrows in (**e**) point to the cilia tip. Scale bar: 2µm. Three biological replicates, n=100 cilia per sample and repetition. **g-h**, KIF7 localisation at the ciliary tip in ciliated NIH3T3 cells treated 3h with solvent control (DMSO) or nocodazole (Noco). Representative cells (**g**) and quantifications (**h**) are shown. The arrows in (**g**) point to the cilia tip. Scale bar: 1.5µm. Three biological replicates, n=100 cilia per sample and repetition. **i**, The percentage of ciliated RPE1 cells with the control or KIF7 siRNA after 48h serum starvation (SS) from Fig. 4e. Three biological replicates, n>100 cilia per sample and repetition. **j**, Percentage of ciliated NIH3T3 cells with the control or KIF7 siRNA after 48h SS from **e**. Three biological replicates, n>100 cilia per sample and repetition. **k-l**, NIH3T3 cells are treated with the indicated siRNA and 48h SS. Representative images (**k**) and quantifications of middle segment (MS, Glu tub) and whole cilia (ARL13B) length (**l**) are shown. The numbers above the cartoons in (**k**) show the ratio of the calculated DS/whole cilia length. Three biological replicates, n=100 cilia per sample and repetition. Scale bar: 2µm. Data shown in (**b, d, f, h, i, j, l)** include mean ±s.d. and *P* values are calculated by two-tailed unpaired student t-test. Source data is shown in Supplementary Table 1.

**Figure S5.**
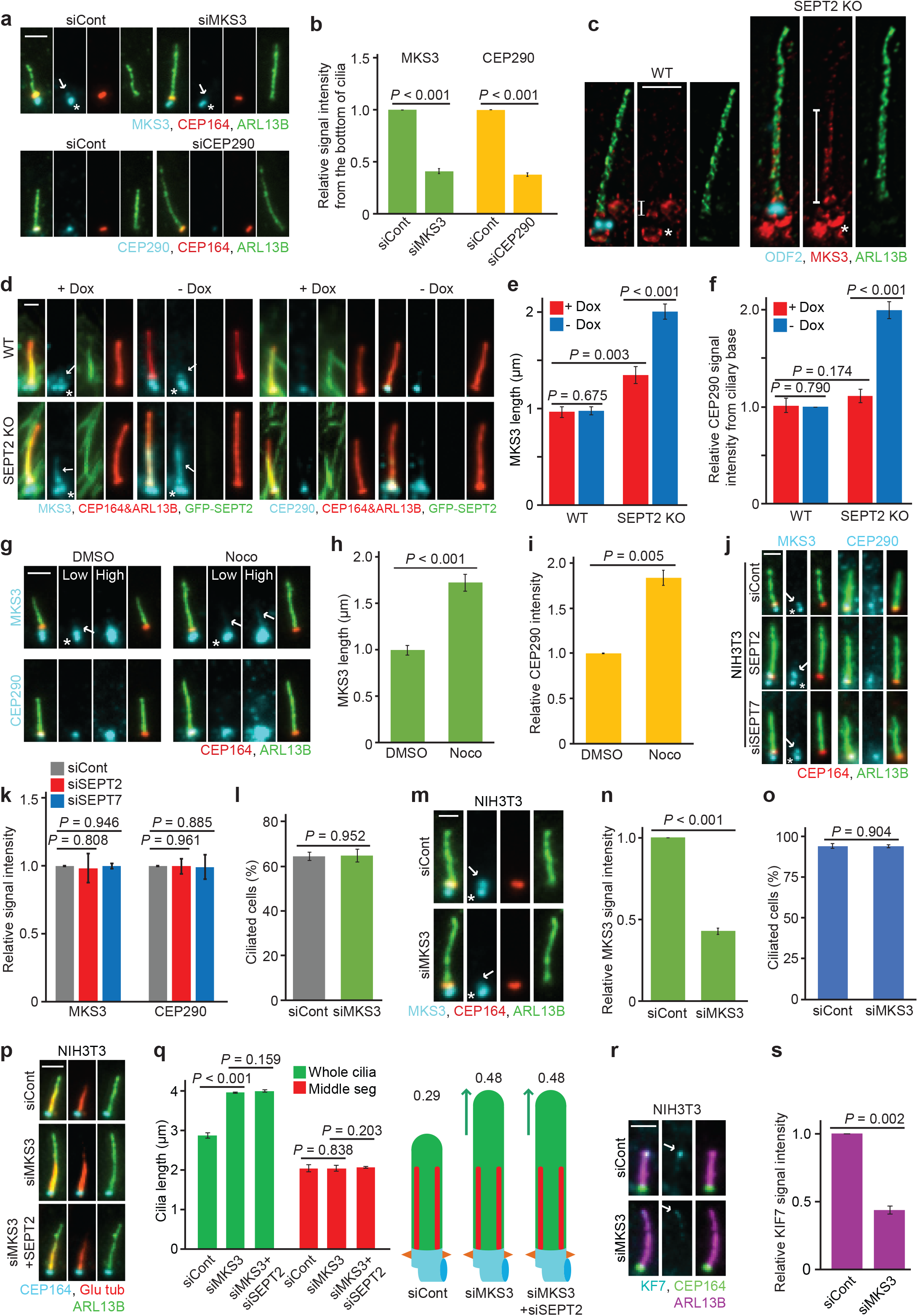
Functional transition zone (TZ) allows to transfer KIF7 to the ciliary tip in RPE1 and NIH3T3 cells. **a-b**, Efficiency of MKS3 and CEP290 knockdown in RPE1 cells. Representative images (**a**) and quantifications of CEP290 or MKS3 protein levels at the TZ (**b**) are shown. White arrows point to MKS3 specific signal and the stars point to unspecific signal that did not decrease upon MKS3 siRNA treatment. Scale bar: 2µm. Three biological replicates, n=100 cilia per sample and replicate. **c**, STED microscopy shows MKS3 cilia localisation in RPE1 WT and SEPT2-KO cells. The white lines indicate the MKS3 specific signal, and the stars point to the unspecific signal. Scale bar: 1µm. **d-f**, Rescue experiment of MKS3 and CEP290 localisation at the cilia after expression of GFP-SEPT2 in RPE1 SEPT2-KO cells. GFP-SEPT2 expression was induced by addition of doxycycline (DOX) as indicated. Representative cells (**d**), MKS3 length at the cilia (**e**) and quantifications of the relative signal intensity of CEP290 at the TZ (**f**) are shown. White arrows point to the specific and stars to unspecific MKS3 signals. Scale bar: 1µm. Three biological replicates, n=100 cilia per sample and repetition. **g**, MKS3 and CEP290 localisation changes in RPE1 ciliated cells treated for 3h with solvent control (DMSO) or nocodazole (Noco). Low and high exposure times are shown. White arrows point to the specific and stars to unspecific MKS3 signal. Scale bar: 2µm. **h**, Quantification of (**g**) showing MKS3 signal length distribution at the cilia. Three biological replicates, n=100 per sample and replicate. **i**, Quantification of (**g**) showing CEP290 relative intensity at the TZ. Three biological replicates, n>90 per sample and replicate. **j**, MKS3 and CEP290 localisation at the TZ in NIH3T3 ciliated cells treated with the indicated siRNAs. White arrows point to the specific and stars to unspecific MKS3 signal. Scale bar: 1.5µm. **k**, Quantification of (**j**) showing normalized MKS3 and CEP290 signal intensities at the TZ. Three biological replicates, n=100 per sample and replicate. **l**, Percentage of ciliated RPE1 cells with the control or MKS3 siRNA after 48h serum starvation (SS) from Fig. 5d. Three biological replicates, n>100 cilia per sample and repetition. **m-n**, Depletion efficiency of MKS3 in ciliated NIH3T3 cells. Representative images (**m**) and quantification (**n**) show MKS3 localisation and signal intensity in ciliated NIH3T3 cells treated with control or MKS3 siRNA. Scale bar: 2µm. Three biological replicates, n>70 per sample and replicate. **o**, Percentage of ciliated NIH3T3 cells with the control or MKS3 siRNA after 48h SS from **m**. Three biological replicates, n>100 cilia per sample and repetition. **p-q**, Influence of MKS3 depletion, but not with SEPT2 co-depletion, on cilia length in NIH3T3 cells. Cells are treated with the indicated siRNA and serum starved (SS) for 48h before immunostaining with the indicated antibodies. Representative images (**p**) and quantifications of middle segment (MS, Glu tub) and whole cilia (ARL13B) length (**q**) are shown. The numbers above the cartoons in (**q**) show the ratio of the calculated DS/whole cilia length. Scale bar: 2µm. Three biological replicates, n=100 cilia per sample and repetition. **r-s**, A functional TZ is required for KIF7 cilia tip localisation in NIH3T3 cells. KIF7 signal intensities are measured in ciliated NIH3T3 cells treated with control or MKS3 siRNA and 48h SS. Representative images (**r**) and quantification of KIF7 signal intensity at the tip of cilia (**s**) are shown. Scale bar: 2µm. Three biological replicates, n=100 cilia per sample and repetition. Data shown in **(b, e, f, h, i, k, l, n, o, q, s)** include mean ±s.d. and *P* values were calculated by two-tailed unpaired student t-test. Source data is shown in Supplementary Table 1.

**Figure S6.**
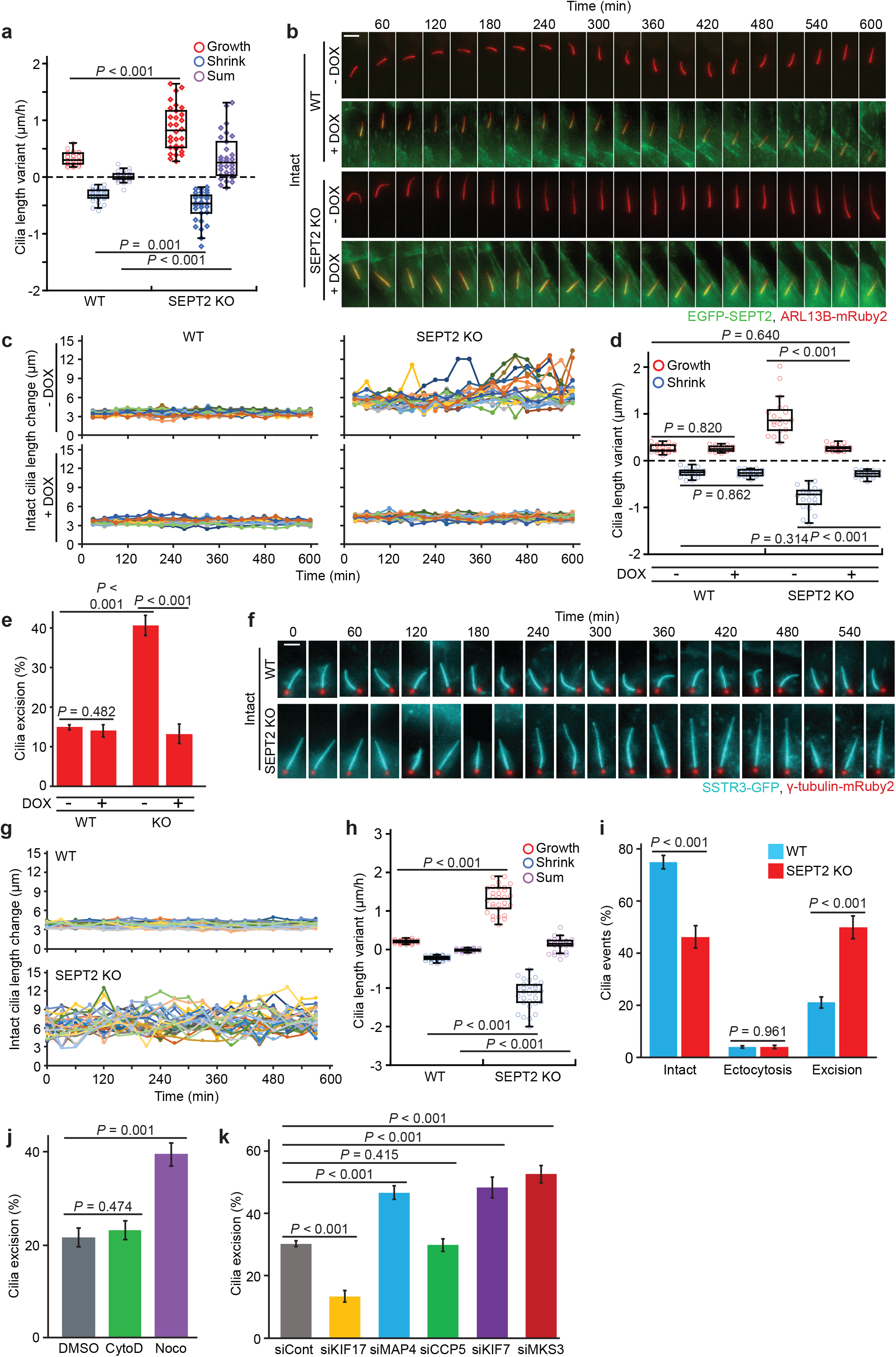
Over-extended distal segments are more prone to undergo excision. **a**, Quantifications show the speed of cilia growth and shortening from Fig. 6a. Red, blue and purple dots show growth, shrink and sum (growth + shrink) speed respectively. n=30 cilia from 3 independent experiments. **b**, Time-lapse images of RPE1 WT and SEPT2-KO cells with (+DOX) or without (-DOX) GFP-SEPT2 expression. Cells are imaged every 30mins for 9.5h. Scale bar: 3µm. Check also **supplementary movie 5-8**. **c**, Quantifications show cilia length variation from (**b**). Two biological replicates, n=10 cilia condition. **d**, Quantifications show the speed of cilia growth and shortening from **b**. Red and blue dots show the speed of growth and shrink, respectively. n=20 cilia from two independent experiments are shown. **e**, Percentage of cilia excision events during live-cell imaging in RPE2 WT and SEPT2-KO RPE1 cells in the presence (+DOX) or absence (-DOX) of GFP-SEPT2. Three biological replicates, n>100 cilia each condition. **f**, Time-lapse images of RPE1 WT and SEPT2-KO cells stably expressing SSTR3-GFP and γ-tubulin-mRuby2. Cells are starved for 32h before the beginning of inspection and imaged every 30min for 9.5h. Scale bar: 2.5µm. For videos, see **supplementary movie 9 and 10**. **g**, Quantification of (**f**) showing changes in cilia length during time-lapse imaging. Three biological replicates, 10 cilia per sample and repetition. **h**, Quantifications show the speed of cilia growth and shortening from (**f**). Red, blue and purple dots show growth, shrink and sum (growth + shrink) speed respectively. n=30 cilia from 3 independent experiments. **i,** Percentage of cilia that remained intact, underwent ectocytosis or excision during live-cell imaging. Three 3 biological replicates, n>100 cilia per sample and repetition. **j**, Percentage of cilia excision events in ciliated RPE1 ARL13B-GFP cells treated with solvent control (DMSO), cytochalasin D (CytoD) or nocodazole (Noco) for 3h. Cilia excision was quantified based on 10h live cell imaging. Three biological replicates, n >100 cilia per sample and repetition. **k**, Percentage of cilia excision events in RPE1 cells treated with the indicated siRNAs. Three biological replicates, Cilia excision was quantified based on 10h live cell imaging. Three biological replicates, n >100 cilia per sample and repetition. Data shown in (**a, d, e, h, i, j, k**) include mean ±s.d.; *P* values are calculated by unpaired Wilcoxon-Mann-Whitney Rank Sum Test (**a, d, h**) or by two-tailed unpaired student t-test (**e, i, j, k**). Source data is shown in Supplementary Table 1.

**Figure S7.**
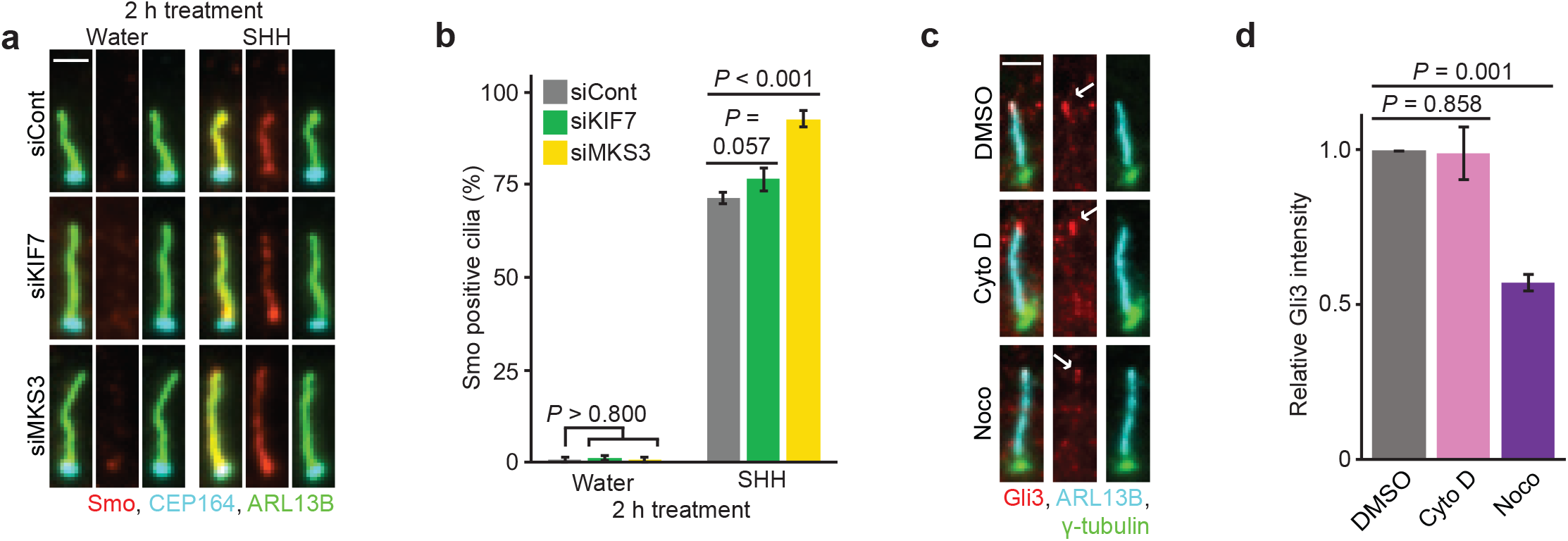
Balancing distal segment length is vital for sonic hedgehog signal transduction. **a-b**, Changes in Smo translocation to the cilium in NIH3T3 cells with indicated siRNA treatment and 48h serum starvation (SS), and then treated with SHH ligands or water for 2h. Smo translocation into the cilium is followed using the indicated antibodies. Representative images (**a**) and quantifications (**b**) are shown. Scale bar: 2µm. Three biological replicates, n>100 cilia per sample and repetition. **c-d**, Gli3 levels at the ciliary tip in ciliated NIH3T3 cells treated with solvent control (DMSO), cytochalasin D (CytoD) or nocodazole (Noco) for 3h. Representative images (**c**) and quantification of Gli3 levels (**d**) are shown. White arrows in (**c**) point to Gli3 at the cilia tip. Scale bar: 2µm. Three biological replicates, n=100 cilia per sample and repetition. Data shown in (**b, d**) include mean ±s.d.; *P* values are calculated by two-tailed unpaired student t-test. Source data are shown in Supplementary Table 1.

**Figure S8.**
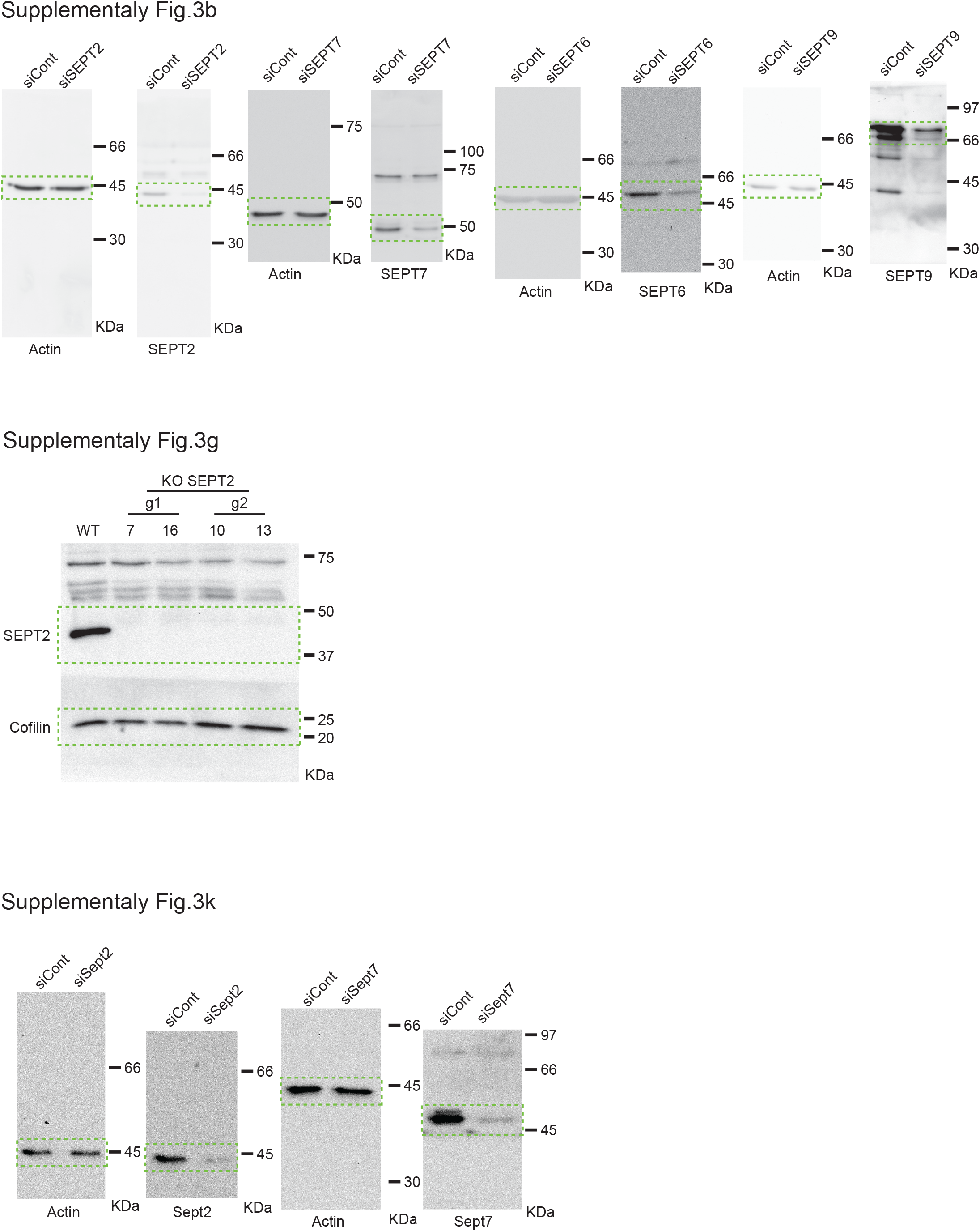
Unprocessed immunoblots. Cropped areas (dashed lines) are shown in supplementary fig. 3b, 3g and 3k as indicated.

## Videos Captions

**Supplementary Movie 1: Cilia dynamics in RPE1 cells.**

Cilia dynamics in RPE1-WT cells stably expressing Arl13b-GFP and γ-tubulin-mRuby2. Live cell imaging was initiated 32-41 h after serum starvation (15 min intervals). The movie corresponds to Figure 6a. Scale bar = 5 µm.

**Supplementary Movie 2: Cilia dynamics in RPE1 SEPT2-KO cells.**

Cilia dynamics in RPE1 SEPT2-KO cells stably expressing Arl13b-GFP and γ-tubulin-mRuby2. Live cell imaging was initiated 32-41 h after serum starvation (15 min intervals). The movie corresponds to Figure 6a. Scale bar = 5 µm.

**Supplementary Movie 3: Ectocytosis in RPE1 SEPT2-KO cells.**

Ectocytosis in RPE1 SEPT2-KO cells stably expressing Arl13b-GFP and γ-tubulin-mRuby2. Live cell imaging was initiated 32-41 h after serum starvation (15 min intervals). The movie corresponds to Figure 6c. Scale bar = 5 µm.

**Supplementary Movie 4: Cilia excision in RPE1 SEPT2-KO cells.**

Cilia excision in RPE1 SEPT2-KO cells stably expressing Arl13b-GFP and γ-tubulin-mRuby2. Live cell imaging was initiated 32-41 h after serum starvation (15 min intervals). The movie corresponds to Figure 6c. Scale bar = 5 µm.

**Supplementary Movie 5: Rescue experiment of cilia growth in RPE1 cells (control cells, no DOX induction).**

Cilia dynamics in RPE1-WT cells stably expressing Arl13b-mRuby2 and EGFP-SEPT2 under control of the doxycycline (DOX) inducible promoter. Live cell imaging was initiated 32-41 h after serum starvation in the absence of DOX (15 min intervals). The movie corresponds to Supplementary Figure 6b. Scale bar = 5 µm.

**Supplementary Movie 6: Rescue experiment of cilia growth in RPE1 cells (control cells, DOX induction).**

Cilia dynamics in RPE1-WT cells stably expressing Arl13b-mRuby2 and EGFP-SEPT2 under control of the doxycycline (DOX) inducible promoter. Cells were serum starved for 32-41 h and 5 ng/ml DOX was added 30 min before starting the movie, for the expression of EGFP-SEPT2 (15 min intervals). The movie corresponds to Supplementary Figure 6b. Scale bar = 5 µm.

**Supplementary Movie 7: Rescue experiment of cilia growth in RPE1 SEPT2-KO cells (control, no DOX induction).**

Cilia dynamics in RPE1 SEPT2-KO cells stably expressing Arl13b-mRuby2 and EGFP-SEPT2 under control of the doxycycline (DOX) inducible promoter. Live cell imaging was initiated 32-41 h after serum starvation in the absence of DOX (15 min intervals). The movie corresponds to Supplementary Figure 6b. Scale bar = 5 µm.

**Supplementary Movie 8: Rescue experiment of cilia growth in RPE1 SEPT2-KO cells with DOX induction.** Cilia dynamics in RPE1 SEPT2-KO cells stably expressing Arl13b-mRuby2 and EGFP-SEPT2 under control of the doxycycline (DOX) inducible promoter. Cells were serum starved for 32-41 h and 5 ng/ml DOX was added 30 min before starting the movie, for the expression of EGFP-SEPT2 (15 min intervals). The movie corresponds to Supplementary Figure 6b. Scale bar = 5 µm.

**Supplementary Movie 9: Cilia dynamics in RPE1 SSTR3-GFP cells.**

Cilia dynamics in RPE1-WT cells stably expressing SSTR3-GFP and γ-tubulin-mRuby2. Live cell imaging was initiated 32-41 h after serum starvation (15 min intervals). The movie corresponds to Supplementary Figure 6f. Scale bar = 5 µm.

**Supplementary Movie 10: Cilia dynamics in RPE1 SEPT2-KO SSTR3-GFP cells.** Cilia dynamics in RPE1 SEPT2-KO cells stably expressing SSTR3-GFP and γ-tubulin-mRuby2. Live cell imaging was initiated 32-41 h after serum starvation (15 min intervals). The movie corresponds to Supplementary Figure 6f. Scale bar = 5 µm.

## References

Aguilar A, Meunier A, Strehl L, Martinovic J, Bonniere M, Attie-Bitach T, Encha-Razavi F & Spassky N (2012) Analysis of human samples reveals impaired SHH-dependent cerebellar development in Joubert syndrome/Meckel syndrome. Proc. Natl. Acad. Sci. U. S. A. 109: 16951–16956 Available at: www.pnas.org/cgi/doi/10.1073/pnas.1201408109 [Accessed March 8, 2021]

Alford LM, Stoddard D, Li JH, Hunter EL, Tritschler D, Bower R, Nicastro D, Porter ME, Sale WS & Street M (2017) The nexin link and B-tubule glutamylation maintain the alignment of outer doublets in the ciliary axoneme. Cytoskeleton 73: 331–340

Baala L, Romano S, Khaddour R, Saunier S, Smith UM, Audollent S, Ozilou C, Faivre L, Laurent N, Foliguet B, Munnich A, Lyonnet S, Salomon R, Encha-Razavi F, Gubler M-C, Boddaert N, de Lonlay P, Johnson CA, Vekemans M, Antignac C, et al (2007) The Meckel-Gruber Syndrome Gene, MKS3, Is Mutated in Joubert Syndrome. Am. J. Hum. Genet. 80: 186–194 Available at: https://www.sciencedirect.com/science/article/pii/S0002929707609331?via#3Dihub [Accessed July 11, 2019]

Besschetnova TY, Kolpakova-Hart E, Guan Y, Zhou J, Olsen BR & Shah J V. (2010) Identification of Signaling Pathways Regulating Primary Cilium Length and Flow-Mediated Adaptation. Curr. Biol. 20: 182–187

van der Burght SN, Rademakers S, Johnson JL, Li C, Kremers GJ, Houtsmuller AB, Leroux MR & Jansen G (2020) Ciliary Tip Signaling Compartment Is Formed and Maintained by Intraflagellar Transport. Curr. Biol. 30: 4299–4306.e5 Available at: https://doi.org/10.1016/j.cub.2020.08.032

Cherry AL, Finta C, Karlström M, Jin Q, Schwend T, Astorga-Wells J, Zubarev RA, Del Campo M, Criswell AR, De Sanctis D, Jovine L & Toftgård R (2013) Structural basis of SUFU-GLI interaction in human Hedgehog signalling regulation. Acta Crystallogr. Sect. D Biol. Crystallogr. 69: 2563–2579

Cheung HOL, Zhang X, Ribeiro A, Mo R, Makino S, Puviindran V, Lo Law KK, Briscoe J & Hui CC (2009) The kinesin protein Kif7 is a critical regulator of Gli transcription factors in mammalian Hedgehog signaling. Sci. Signal. 2: ra29– ra29 Available at: https://stke.sciencemag.org/content/2/76/ra29 [Accessed July 31, 2020]

Chih B, Liu P, Chinn Y, Chalouni C, Komuves LG, Hass PE, Sandoval W & Peterson AS (2012) A ciliopathy complex at the transition zone protects the cilia as a privileged membrane domain. Nat. Cell Biol. 14: 61–72 Available at: https://pubmed.ncbi.nlm.nih.gov/22179047/ [Accessed January 15, 2021]

Cornelia I. Bargmann (2006) Chemosensation in C. elegans.

Echelard Y, Epstein DJ, St-Jacques B, Shen L, Mohler J, McMahon JA & McMahon AP (1993) Sonic hedgehog, a member of a family of putative signaling molecules, is implicated in the regulation of CNS polarity. Cell 75: 1417–1430

Endoh-Yamagami S, Evangelista M, Wilson D, Wen X, Theunissen JW, Phamluong K, Davis M, Scales SJ, Solloway MJ, de Sauvage FJ & Peterson AS (2009) The Mammalian Cos2 Homolog Kif7 Plays an Essential Role in Modulating Hh Signal Transduction during Development. Curr. Biol. 19: 1320–1326

Flood PR & Totland GK (1977) Substructure of solitary cilia in mouse kidney. Cell Tissue Res. 183: 281–290

Gadadhar S, Dadi H, Bodakuntla S, Schnitzler A, Bièche I, Rusconi F & Janke C (2017) Tubulin glycylation controls primary cilia length. J. Cell Biol. 216: 2701– 2713

Garcia-Gonzalo FR & Reiter JF (2017) Open Sesame: How Transition Fibers and the Transition Zone Control Ciliary Composition. Cold Spring Harb. Perspect. Biol. 9: a028134 Available at: http://www.ncbi.nlm.nih.gov/pubmed/27770015 [Accessed April 11, 2019]

Ghossoub R, Hu Q, Failler M, Rouyez M-C, Spitzbarth B, Mostowy S, Wolfrum U, Saunier S, Cossart P, James Nelson W & Benmerah A (2013) Septins 2, 7 and 9 and MAP4 colocalize along the axoneme in the primary cilium and control ciliary length. J. Cell Sci. 126: 2583–2594 Available at: http://jcs.biologists.org/cgi/doi/10.1242/jcs.111377

Guen VJ, Gamble C, Perez DE, Bourassa S, Zappel H, Gärtner J, Lees JA & Colas P (2016) STAR syndrome-associated CDK10/Cyclin M regulates actin network architecture and ciliogenesis. Cell Cycle 15: 678–88 Available at: http://www.ncbi.nlm.nih.gov/pubmed/27104747 [Accessed January 3, 2018]

Guo S, Liao H, Liu J, Liu J, Tang F, He Z, Li Y & Yang Q (2018) Resveratrol Activated Sonic Hedgehog Signaling to Enhance Viability of NIH3T3 Cells ***in Vitro*** via Regulation of Sirt1. Cell. Physiol. Biochem. 50: 1346–1360 Available at: https://www.karger.com/Article/FullText/494593 [Accessed June 25, 2020]

Hao L, Thein M, Brust-Mascher I, Civelekoglu-Scholey G, Lu Y, Acar S, Prevo B, Shaham S & Scholey JM (2011) Intraflagellar transport delivers tubulin isotypes to sensory cilium middle and distal segments. Nat. Cell Biol. 13: 790–8 Available at: http://www.ncbi.nlm.nih.gov/pubmed/21642982 [Accessed February 13, 2019]

Hata S, Pastor Peidro A, Panic M, Liu P, Atorino E, Funaya C, Jäkle U, Pereira G & Schiebel E (2019) The balance between KIFC3 and EG5 tetrameric kinesins controls the onset of mitotic spindle assembly. Nat. Cell Biol. 21: 1138–1151 Available at: http://www.nature.com/articles/s41556-019-0382-6 [Accessed September 17, 2019]

Haycraft CJ, Banizs B, Aydin-Son Y, Zhang Q, Michaud EJ & Yoder BK (2005) Gli2 and Gli3 Localize to Cilia and Require the Intraflagellar Transport Protein Polaris for Processing and Function. Available at: www.plosgenetics.org [Accessed January 28, 2019]

He K, Ling K & Hu J (2020) The emerging role of tubulin posttranslational modifications in cilia and ciliopathies. Biophys. Reports 6: 89–104 Available at: https://doi.org/10.1007/s41048-020-00111-0 [Accessed October 5, 2020]

He K, Ma X, Xu T, Li Y, Hodge A, Zhang Q, Torline J, Huang Y, Zhao J, Ling K & Hu J (2018) Axoneme polyglutamylation regulated by Joubert syndrome protein ARL13B controls ciliary targeting of signaling molecules. Nat. Commun. 9: 3310 Available at: http://www.nature.com/articles/s41467-018-05867-1 [Accessed September 26, 2018]

He M, Subramanian R, Bangs F, Omelchenko T, Liem KF, Kapoor TM & Anderson K V. (2014) The kinesin-4 protein Kif7 regulates mammalian Hedgehog signalling by organizing the cilium tip compartment. Nat. Cell Biol. 16: 663–672

Hu Q, Milenkovic L, Jin H, Scott MP, Nachury M V, Spiliotis ET & Nelson WJ (2010) A Septin Diffusion Barrier at the Base Membrane Protein Distribution. Science (80-.). 329: 436–439

Insinna C, Pathak N, Perkins B, Drummond I & Besharse JC (2008) The homodimeric kinesin, Kif17, is essential for vertebrate photoreceptor sensory outer segment development. Dev. Biol. 316: 160–170 Available at: https://pubmed.ncbi.nlm.nih.gov/18304522/ [Accessed July 27, 2020]

Ishikawa H & Marshall WF (2011) Ciliogenesis: building the cell’s antenna. Nat. Rev. Mol. Cell Biol. 12: 222–234 Available at: http://www.nature.com/articles/nrm3085 [Accessed November 12, 2018]

Kim J, Lee JE, Heynen-Genel S, Suyama E, Ono K, Lee K, Ideker T, Aza-Blanc P & Gleeson JG (2010) Functional genomic screen for modulators of ciliogenesis and cilium length. Nature 464: 1048–1051 Available at: http://www.nature.com/doifinder/10.1038/nature08895 [Accessed January 8, 2018]

Kim S & Dynlacht BD (2013) Assembling a primary cilium. Curr. Opin. Cell Biol. 25: 506–511

Kim Y, Osborn DP, Lee J, Araki M, Araki K, Mohun T, Känsäkoski J, Brandstack N, Kim HH, Miralles F, Kim C, Brown NA, Kim HH, Martinez-Barbera JP, Ataliotis P, Raivio T, Layman LC & Kim S (2018) WDR11-mediated Hedgehog signalling defects underlie a new ciliopathy related to Kallmann syndrome. EMBO Rep. 19: 269–289 Available at: https://onlinelibrary.wiley.com/doi/10.15252/embr.201744632 [Accessed March 30, 2021]

Kramer JM, Moerman DG & Inglis PN (2007) The sensory cilia of Caenorhabditis elegans. Available at: http://www.wormbook.org. [Accessed November 15, 2018]

Lechtreck KF & Geimer S (2000) Distribution of polyglutamylated tubulin in the flagellar apparatus of green flagellates. Cell Motil. Cytoskeleton 47: 219–235 Available at: https://onlinelibrary.wiley.com/doi/epdf/10.1002/1097-0169%28200011%2947%3A3%3C219%3A%3AAID-CM5%3E3.0.CO%3B2-Q [Accessed July 7, 2020]

Lee J, Chung YD & Doo Chung Y (2015) Ciliary subcompartments: how are they established and what are their functions? BMB Rep 48: 380–387 Available at: www.bmbreports.org http://dx.doi.org/10.5483/BMBRep.2015.48.7.084 [Accessed January 31, 2019]

Leightner AC, Hommerding CJ, Peng Y, Salisbury JL, Gainullin VG, Czarnecki PG, Sussman CR & Harris PC (2013) The Meckel syndrome protein meckelin (TMEM67) is a key regulator of cilia function but is not required for tissue planar polarity. Hum. Mol. Genet. 22: 2024–2040 Available at: https://academic.oup.com/hmg/article-lookup/doi/10.1093/hmg/ddt054 [Accessed April 10, 2019]

Lewis TR, Kundinger SR, Pavlovich AL, Bostrom JR, Link BA & Besharse JC (2017) Cos2/Kif7 and Osm-3/Kif17 regulate onset of outer segment development in zebrafish photoreceptors through distinct mechanisms. Dev. Biol. 425: 176–190

Li ZJ, Nieuwenhuis E, Nien W, Zhang X, Zhang J, Puviindran V, Wainwright BJ, Kim PCW & Hui C chung (2012) Kif7 regulates Gli2 through Sufu-dependent and -independent functions during skin development and tumorigenesis. Dev. 139: 4152–4161 Available at: https://dev.biologists.org/content/139/22/4152 [Accessed July 31, 2020]

May-Simera HL, Wan Q, Jha BS, Hartford J, Khristov V, Dejene R, Chang J, Patnaik S, Lu Q, Banerjee P, Silver J, Insinna-Kettenhofen C, Patel D, Lotfi M, Malicdan M, Hotaling N, Maminishkis A, Sridharan R, Brooks B, Miyagishima K, et al (2018) Primary Cilium-Mediated Retinal Pigment Epithelium Maturation Is Disrupted in Ciliopathy Patient Cells. Cell Rep. 22: 189–205 Available at: https://doi.org/10.1016/j.celrep.2017.12.038 [Accessed January 13, 2020]

Miyoshi K, Kasahara K, Murakami S, Takeshima M, Kumamoto N, Sato A, Miyazaki I, Matsuzaki S, Sasaoka T, Katayama T & Asanuma M (2014) Lack of Dopaminergic Inputs Elongates the Primary Cilia of Striatal Neurons. PLoS One 9: e97918 Available at: https://dx.plos.org/10.1371/journal.pone.0097918 [Accessed January 6, 2020]

Mönnich M, Borgeskov L, Breslin L, Jakobsen L, Rogowski M, Doganli C, Schrøder JM, Mogensen JB, Blinkenkjær L, Harder LM, Lundberg E, Geimer S, Christensen ST, Andersen JS, Larsen LA & Pedersen LB (2018) CEP128 Localizes to the Subdistal Appendages of the Mother Centriole and Regulates TGF-β/BMP Signaling at the Primary Cilium. Cell Rep. 22: 2584–2592 Available at: http://www.ncbi.nlm.nih.gov/pubmed/29514088 [Accessed March 28, 2018]

Mukhopadhyay S, Lu Y, Shaham S & Sengupta P (2008) Sensory Signaling-Dependent Remodeling of Olfactory Cilia Architecture in C. elegans. Dev. Cell 14: 762–774 Available at: https://pubmed.ncbi.nlm.nih.gov/18477458/ [Accessed February 25, 2021]

Nachury M V. (2014) How do cilia organize signalling cascades? Philos. Trans. R. Soc. B Biol. Sci. 369: 20130465–20130465 Available at: http://rstb.royalsocietypublishing.org/cgi/doi/10.1098/rstb.2013.0465 [Accessed November 15, 2018]

Nager AR, Goldstein JS, Herranz-Pérez V, Portran D, Ye F, Garcia-Verdugo JM & Nachury M V. (2017) An Actin Network Dispatches Ciliary GPCRs into Extracellular Vesicles to Modulate Signaling. Cell 168: 252–263.e14

Neubauer K & Zieger B (2017) The Mammalian Septin Interactome. Front. Cell Dev. Biol. 5: 1–9 Available at: http://journal.frontiersin.org/article/10.3389/fcell.2017.00003/full

O’Hagan R, Piasecki BP, Silva M, Phirke P, Nguyen KCQ, Hall DH, Swoboda P & Barr MM (2011) The tubulin deglutamylase CCPP-1 regulates the function and stability of sensory cilia in C. elegans. Curr. Biol. 21: 1685–1694 Available at: /pmc/articles/PMC4680987/?report=abstract [Accessed November 28, 2019]

Oh EC & Katsanis N (2013) Context-dependent regulation of Wnt signaling through the primary cilium. J. Am. Soc. Nephrol. 24: 10–18 Available at: www.jasn.org [Accessed March 30, 2021]

Palander O, El-Zeiry M & Trimble WS (2017) Uncovering the Roles of Septins in Cilia. Front. Cell Dev. Biol. 5: 1–7 Available at: http://journal.frontiersin.org/article/10.3389/fcell.2017.00036/full

Pedersen LB & Akhmanova A (2014) Kif7 keeps cilia tips in shape. Nat. Cell Biol. 16: 623–625

Pedersen LB, Mogensen JB & Christensen ST (2016) Endocytic Control of Cellular Signaling at the Primary Cilium. Trends Biochem. Sci. 41: 784–797

Prevo B, Scholey JM & Peterman EJG (2017) Intraflagellar Transport: Mechanisms of Motor Action, Cooperation and Cargo Delivery. FEBS J. 284: 2905–2931 Available at: http://doi.wiley.com/10.1111/febs.14068 [Accessed February 13, 2019]

Qiu N, Xiao Z, Cao L, Buechel MM, David V, Roan E & Quarles LD (2012) Disruption of Kif3a in osteoblasts results in defective bone formation and osteopenia. J. Cell Sci. 125: 1945–1957

Ramsbottom SA, Molinari E, Srivastava S, Silberman F, Henry C, Alkanderi S, Devlin LA, White K, Steel DH, Saunier S, Miles CG & Sayer JA (2018) Targeted exon skipping of a CEP290 mutation rescues Joubert syndrome phenotypes in vitro and in a murine model. Proc. Natl. Acad. Sci. 115: 12489–12494 Available at: https://www.pnas.org/content/115/49/12489 [Accessed July 12, 2019]

Reiter JF & Leroux MR (2017) Genes and molecular pathways underpinning ciliopathies. Nat. Rev. Mol. Cell Biol. 18: 533–547 Available at: http://dx.doi.org/10.1038/nrm.2017.60 [Accessed January 2, 2018]

Sánchez I & Dynlacht BD (2016) Cilium assembly and disassembly. Nat. Cell Biol. 18: 711–717 Available at: http://www.nature.com/articles/ncb3370 [Accessed November 3, 2018]

Sandrock K, Bartsch I, Bläser S, Busse A, Busse E & Zieger B (2011) Characterization of human septin interactions. Biol. Chem. 392: 751–761 Available at: https://www.degruyter.com/view/j/bchm.2011.392.issue-8-9/bc.2011.081/bc.2011.081.xml [Accessed January 4, 2018]

Schindelin J, Arganda-Carreras I, Frise E, Kaynig V, Longair M, Pietzsch T, Preibisch S, Rueden C, Saalfeld S, Schmid B, Tinevez JY, White DJ, Hartenstein V, Eliceiri K, Tomancak P & Cardona A (2012) Fiji: An open-source platform for biological-image analysis. Nat. Methods 9: 676–682 Available at: http://fiji.sc/Adding_Update_Sites [Accessed May 6, 2021]

Sharma N, Kosan ZA, Stallworth JE, Berbari NF & Yoder BK (2011) Soluble levels of cytosolic tubulin regulate ciliary length control. Available at: http://www.molbiolcell.org/cgi/ [Accessed November 15, 2018]

Shi X, Garcia G, Van De Weghe JC, McGorty R, Pazour GJ, Doherty D, Huang B & Reiter JF (2017) Super-resolution microscopy reveals that disruption of ciliary transition-zone architecture causes Joubert syndrome. Nat. Cell Biol. 19: 1178– 1188 Available at: http://www.nature.com/doifinder/10.1038/ncb3599 [Accessed March 13, 2018]

Shi X, Zhan X & Wu J (2015) A positive feedback loop between Gli1 and tyrosine kinase Hck amplifies shh signaling activities in medulloblastoma. Oncogenesis 4:

Slaats GG, Isabella CR, Kroes HY, Dempsey JC, Gremmels H, Monroe GR, Phelps IG, Duran KJ, Adkins J, Kumar SA, Knutzen DM, Knoers N V, Mendelsohn NJ, Neubauer D, Mastroyianni SD, Vogt J, Worgan L, Karp N, Bowdin S, Glass IA, et al (2016) MKS1 regulates ciliary INPP5E levels in Joubert syndrome. J. Med. Genet. 53: 62–72 Available at: http://www.ncbi.nlm.nih.gov/pubmed/26490104 [Accessed July 12, 2019]

Snow JJ, Ou G, Gunnarson AL, Regina M, Walker S, Zhou HM, Brust-Mascher I & Scholey JM (2004a) Two Anterograde Intraflagellar Transport Motors Cooperate to Build Distinct Parts of Sensory Cilia on Caenorhabditis elegans Neurons Available at: https://cloudfront.escholarship.org/dist/prd/content/qt6737m19b/qt6737m19b.pdf?t=lnqgql [Accessed February 13, 2019]

Snow JJ, Ou G, Gunnarson AL, Walker MRS, Zhou HM, Brust-Mascher I, Scholey JM, Regina M, Walker S, Zhou HM, Brust-Mascher I & Scholey JM (2004b) Two anterograde intraflagellar transport motors cooperate to build sensory cilia on C. elegans neurons. Nat. Cell Biol. 6: 1109–1113 Available at: https://www.nature.com/articles/ncb1186.pdf [Accessed February 13, 2019]

Srivastava S, Ramsbottom SA, Molinari E, Alkanderi S, Filby A, White K, Henry C, Saunier S, Miles CG & Sayer JA (2017) A human patient-derived cellular model of Joubert syndrome reveals ciliary defects which can be rescued with targeted therapies. Hum. Mol. Genet. 26: 4657–4667 Available at: https://pdfs.semanticscholar.org/e2d8/fabdda29b783e0aa8dd656a6c4d4bd43630a.pdf [Accessed July 11, 2019]

Sun S, Fisher RL, Bowser SS, Pentecost BT & Sui H (2019) Three-dimensional architecture of epithelial primary cilia. Proc. Natl. Acad. Sci. U. S. A. 116: 9370– 9379

Sun X, Park JH, Gumerson J, Wu Z, Swaroop A, Qian H, Roll-Mecak A & Li T (2016) Loss of RPGR glutamylation underlies the pathogenic mechanism of retinal dystrophy caused by TTLL5 mutations. Proc. Natl. Acad. Sci. U. S. A. 113: E2925–E2934 Available at: www.pnas.org/cgi/doi/10.1073/pnas.1523201113 [Accessed January 7, 2021]

Tammachote R, Hommerding CJ, Sinders RM, Miller CA, Czarnecki PG, Leightner AC, Salisbury JL, Ward CJ, Torres VE, Gattone VH & Harris PC (2009) Ciliary and centrosomal defects associated with mutation and depletion of the Meckel syndrome genes MKS1 and MKS3. Hum. Mol. Genet. 18: 3311–3323 Available at: https://academic.oup.com/hmg/article-lookup/doi/10.1093/hmg/ddp272 [Accessed July 12, 2019]

Taylor SP, Dantas TJ, Duran I, Wu S, Lachman RS, Nelson SF, Cohn DH, Vallee RB, Krakow D, Nelson SF, Cohn DH, Vallee RB & Krakow D (2015) Mutations in DYNC2LI1 disrupt cilia function and cause short rib polydactyly syndrome. Nat. Commun. 6: 7092 Available at: http://www.nature.com/articles/ncomms8092 [Accessed February 13, 2019]

Valadares NF, d’Muniz Pereira H, Ulian Araujo AP & Garratt RC (2017) Septin structure and filament assembly. Biophys. Rev. 9: 481–500 Available at: http://www.ncbi.nlm.nih.gov/pubmed/28905266 [Accessed January 10, 2018]

Wann AKT & Knight MM (2012) Primary cilia elongation in response to interleukin-1 mediates the inflammatory response. Cell. Mol. Life Sci. 69: 2967–2977

Waters AM & Beales PL (2011) Ciliopathies: an expanding disease spectrum. Pediatr. Nephrol. 26: 1039–1056 Available at: http://link.springer.com/10.1007/s00467-010-1731-7 [Accessed January 13, 2020]

Wheway G, Nazlamova L & Hancock JT (2018) Signaling through the Primary Cilium. Front. Cell Dev. Biol. 6: 8 Available at: http://journal.frontiersin.org/article/10.3389/fcell.2018.00008/full [Accessed January 16, 2019]

Wloga D, Joachimiak E, Louka P & Gaertig J (2017) Posttranslational modifications of Tubulin and cilia. Cold Spring Harb. Perspect. Biol. 9: Available at: http://cshperspectives.cshlp.org/ [Accessed November 28, 2019]

Wood CR, Huang K, Diener DR & Rosenbaum JL (2013) The cilium secretes bioactive ectosomes. Curr. Biol. 23: 906–911 Available at: https://pubmed.ncbi.nlm.nih.gov/23623554/ [Accessed May 13, 2021]

